# Computational Analysis of Therapeutic Neuroadaptation to Chronic Antidepressant in a Model of the Monoaminergic Neurotransmitter, Stress Hormone, and Male Sex Hormone Systems

**DOI:** 10.1101/634030

**Authors:** Mariam Bonyadi Camacho, Warut D. Vijitbenjaronk, Thomas J Anastasio

**Affiliations:** Computational Neurobiology Laboratory, Neuroscience Program, Medical Scholars Program, University of Illinois College of Medicine at Urbana-Champaign, Urbana, IL, USA; Computational Neurobiology Laboratory, Department of Computer Science, University of Illinois at Urbana-Champaign, Urbana, IL, USA; Computational Neurobiology Laboratory, Department of Molecular and Integrative Physiology, University of Illinois at Urbana-Champaign, Urbana, IL, USA

**Keywords:** neuropharmacology, drug discovery, machine learning, systems biology, gonadal hormones, depression

## Abstract

Second-line depression treatment involves augmentation with one (rarely two) additional drugs, of chronic administration of a selective serotonin reuptake inhibitor (SSRI), which is the first-line depression treatment. Unfortunately, many depressed patients still fail to respond even after months to years of searching to find an effective combination. To aid in the identification of potentially affective antidepressant combinations, we created a computational model of the monoaminergic neurotransmitter (serotonin, norepinephrine, and dopamine), stress-hormone (cortisol), and male sex-hormone (testosterone) systems. The model was trained via machine learning to represent a broad set of empirical observations. Neuroadaptation to chronic drug administration was simulated through incremental adjustments in model parameters that corresponded to key regulatory components of the neurotransmitter and neurohormone systems. Analysis revealed that neuroadaptation in the model depended on all of the regulatory components in complicated ways, and did not reveal any one or a few specific components that could be targeted in the design of combination antidepressant treatment. We used large sets of neuroadapted states of the model to screen 74 different drug and hormone combinations and identified several combinations that could potentially be therapeutic for a higher proportion of male patients than SSRIs by themselves.

## 1. Introduction

The majority of antidepressant drugs in use today were derived on the basis of the monoamine hypothesis, which originated from clinical observations in the 1960s that drugs that elevate brain monoamine levels also elevate mood (Schildkraut, 1965). The monoamines are: serotonin (5HT), secreted by dorsal raphe (DR); norepinephrine (NE), secreted by locus coeruleus (LC); and dopamine (DA), secreted by ventral tegmental area (VTA). Due to their more tolerable side-effect profiles, chronic administration of selective serotonin reuptake inhibitors (SSRIs) has now replaced chronic administration of other monoamine-targeting drugs as the first-line pharmacological intervention for depression (Koenig and Thase, 2009). Unfortunately, chronic SSRI is only about 30% effective (Cipriani et al., 2016; Rush et al., 2006; Trivedi et al., 2006a; Turner et al., 2008). Combining chronic SSRI with another monoamine-targeting drug can raise efficacy, but finding an effective combination can take months or years and still fail to provide relief for the 10-30% of depressed patients classified as treatment-resistant (Ananth, 1998; Barowsky and Schwartz, 2006; Trivedi et al., 2006b; Zhou et al., 2015). The drawbacks associated with monoamine-targeting drugs has motivated the depression field to investigate the relationships between the monoaminergic and the neuroendocrine systems to identify new pharmacological strategies for depression treatment (Berton and Nestler, 2006).

To aid the identification of potentially effective depression polytherapies, we developed a computational model of the neuroendocrine system that represents many of the known interactions between the monoaminergic nervous system and the stress- and sex-hormone systems. The monoamine-stress-sex model (MSS-model) is an extension of our monoamine-stress model (MS-model), which is itself an extension of our monoamine model (M-model) (Camacho and Anastasio, 2017; Camacho et al., 2018). The MSS-model represents interactions between the 3 monoaminergic neurotransmitter systems (centered on DR, LC, and VTA), the neuroendocrine systems that regulate cortisol (CORT, centered on the paraventricular nucleus (PVN) and hypothalamic-pituitary-adrenal (HPA) axis), and testosterone (TEST, centered on the preoptic area (POA) and the hypothalamic-pituitary-gonadal (HPG) axis) (the MSS-model is focused on the male neuroendocrine system; see Discussion).

Chronic (days to weeks) antidepressant administration is known to cause adaptive changes in neurons within the DR, LC, and VTA that bring their activity levels back toward their normative, pre-drug, baseline levels of activity (Blier and De Montigny, 1987; De Montigny et al., 1990). The MSS-model simulates adaptive changes by allowing a subset of transmitter-system components (TSCs, which are proteins such as receptors and transporters that set neurotransmitter or neurohormone tone) to increase or decrease the strength of their influence (representing changes in protein expression, sensitivity, or cellular/synaptic localization) on other elements in the network. We refer to TSC-strength configurations that restore DR, LC, and VTA activity back toward baseline in the MSS-model as “adapted.” Adapted TSC-strength configurations were considered “therapeutic” if they caused the MSS-model to elevate the monoamines and reduce cortisol as described in clinical studies of responders to antidepressant therapy (Ceglia et al., 2004; Ruhé et al., 2015; Schildkraut, 1965).

Through various forms of computational analysis (i.e. sensitivity, correlation, temporal-logic, and state-space), we show that no single TSC or pair of TSCs determine therapeutic monoamine/cortisol levels by themselves, but that achievement of a therapeutic outcome involves all of the TSCs interacting in a complex way. This complexity essentially precludes design of polytherapies that target specific TSCs but further justifies the use of the MSS-model, which represents those complex monoamine/neuroendocrine interactions, as a computational screen to identify novel drug/hormone combinations. By generating a large set of TSC-strength configurations adapted to SSRI and each of 74 different chronic drug/hormone combinations in a computational model of the male MSS system, we identified candidate combinations that could potentially provide symptomatic relief to a larger proportion of depressed male patients than the current first-line approach. Additionally, our modeling results support preliminary clinical findings that augmentation of SSRI action with Testosterone can provide depression relief in a larger proportion of male depressives (Kanayama et al., 2007; Scantamburlo et al., 2015).

## 2. Experimental procedures

### 2.1 MSS-model Formalism Overview

The MSS-model has a “foreground” and a “background.” The foreground is a representation of the main components of the monoaminergic-transmitter, stress- and sex-hormone systems. The background is a representation of most of the related neural, transmitter, and hormonal systems that are known to interact with the monoamine-stress-sex systems but on which fewer data are available. Single elements of the MSS-model represented key brain regions including all of the foreground brain regions (e.g. DR, LC, VTA, PVN, and POA) and key molecular components including receptors, transporters, enzymes, precursors, metabolites, and the transmitters and hormones themselves. See Supplemental Material for a complete listing of abbreviations.

The MSS-model was constructed, parameterized, run, adapted, and analyzed in the same way as the MS-model (for details see Camacho et al., 2018). Computational and analytical procedures will be briefly summarized here and any differences with the MS-model will be noted.

The MSS-model takes the form of a recurrent network of nonlinear elements or “units”. All of the units have the same sigmoidal (S-shaped) nonlinearity that bounds their activity between 0 and 1 and produces an output of 0.50 for 0 input. Units in the network are classified as input, output, or “hidden”. Input units send connections to hidden and output units, while hidden and output units send connections to each other but not to input units. Input values are preassigned to input units, and desired (or target) values are preassigned to output units during training. Hidden units have no preassigned values. All connections in the network are weighted. On each network update, each hidden or output unit computes the sum of its weighted inputs from other units and applies the sigmoid to its weighted input sum.

The units in the model correspond to regional or brain-wide biological variables. Some single units represented the average activity of all of the neurons in specific brain regions while others represented brain levels of key transmitters and hormones. Still others represented the overall brain “activity” (expression, sensitivity, concentration, etc., or combinations thereof) of key receptors, transporters, enzymes, and small molecules that constitute transmitter-system components (TSCs). While the units represented regional or brain-wide variables, the weights of the connections between units represented cellular-level variables. The effectiveness of the 5HT transporter (5HTT) in inhibiting 5HT levels, for example, was represented in the model by the absolute value of the inhibitory weight of the 5HTT unit onto the 5HT unit. Figure 1 shows a highly simplified model diagram. The full model diagram can be interactively viewed in Supplemental Figure 1.

**Figure 1:**
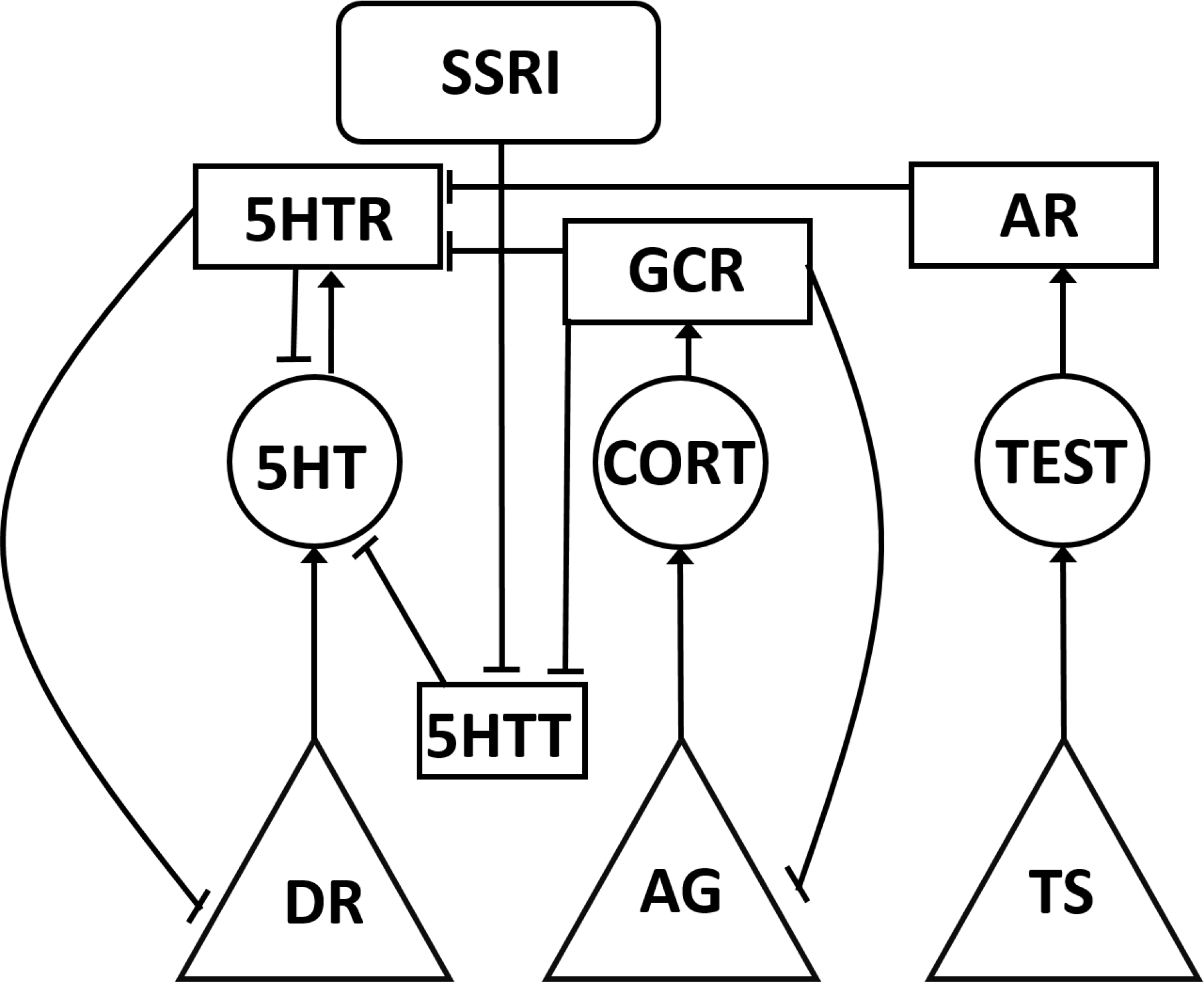
Simplified model diagram. The monoamine-stress-sex model (MSS-model) takes the form of a recurrent network of non-linear elements (units) representing neurotransmitter and hormone producing brain regions, as well as key molecular species including enzymes, transmitters, hormones, and receptors. Each unit type is represented in this schematic representation using a different shape. Only one or two of each unit type from each system (monoamine, stress, or sex system) is shown for simplicity. Neurotransmitter and hormone producing regions, neurotransmitters and hormones, protein molecules, or inputs are represented respectively as triangles, circles, rectangles, or rounded rectangles. Connections between model units are represented with arrows or tees, representing excitatory and inhibitory connections, respectively.

A small subset (precisely 13) of the connection weights represent cell-specific entities that are known to adapt or adjust their strengths (through changes in expression, sensitivity, concentration, etc.) under chronic stress, or with chronic administration of drugs or hormones. These will be referred to as “adjustable TSCs”. The adjustable-TSC weights were used to analyze adaptation (specifically, neuroadaptation) to chronic administration of drugs, hormones, and combinations in the MSS-model.

### 2.2 Model Structure and Function

There were 132 total network units (54 input, 32 output, and 46 hidden units). The 77×130 network weight matrix organizes the connection weights from the input, hidden, and output units to the hidden and output units for a total of 10,010 connection weights. There are 3 major classes of connection weights in the MSS-model: canonical, structure, and non-structure.

The 36 canonical weights (listed in Supplemental Table 2) represent empirically known interactions between key elements of the foreground systems: monoaminergic system, HPA axis, and HPG axis. One canonical weight, for example, is the weight of the connection from the testes unit to the TEST unit. This canonical weight represents the efficacy of the testes in releasing testosterone. The 426 structure weights represent other empirically known interactions between elements of the background systems, such as the connection weight between the LC and galanin units, representing the efficacy of galanin co-released by the LC. The 9,548 non-structure weights are all the other connection weights which represent interactions that have not been identified yet and may or may not represent interactions that occur neurobiologically.

The MSS-model was constructed and parameterized on the basis of two data structures: a structure matrix and a truth table, both of which were populated through extensive literature review. The structure matrix is a 77×130 matrix that is coextensive with the network weight matrix, which specifies the canonical and structure connections between model units as well as their valence (positive or negative) if known (see Supplemental Material S1).

The truth table is an array of input/desired-output training patterns that specifies how specific inputs (experimental manipulations such as drugs, stress, hormones, lesions, etc.) are known to affect specific output units (such as 5HT, CORT, or TEST level). Each row of the truth-table represents the results of at least one experiment. Inputs can be either present (1) or absent (0). Desired outputs specify maximal decrease, moderate decrease, no change, moderate increase, or maximal increase, corresponding to values of 0.30, 0.40, 0.50, 0.60, and 0.70, respectively (see Supplemental Material S2). Table 1 is a condensed, sample input/desired-output table. The complete truth-table can be viewed in Supplemental Table 1.

**Table 1:** Highly simplified example of truth-table data used to train the Monoamine-Stress-Sex model.

| Input | Desired Output |  |  |  |
| --- | --- | --- | --- | --- |
|  | DR | Testosterone<sub>end</sub> | 5HT | CORT |
| Baseline | 0.50 | 0.60 | 0.50 | 0.50 |
| SSRI | 0.40 | 0.60 | 0.60 | 0.70 |
| Stress | 0.60 | 0.50 | 0.60 | 0.70 |
| Testosterone<sub>ex</sub> | 0.60 | 0.70 | 0.60 |  |
| Stress+Castration |  |  | 0.60 | 0.70 |

### 2.3 Training the Model

Model parameterization involved training its connection weights using a recurrent back-propagation algorithm which trains dynamic, recurrent neural networks to produce desired steady-state outputs in response to specific inputs (Pineda, 1987). Model training involved two main sources of randomness. The entire connection weight matrix was randomized prior to training, and then the network was trained by presenting input/desired-output patterns selected at random, repeatedly and with replacement, over 1×10^6^ training cycles. Networks trained from different random starting weight matrices and according to different random orders of input/desired-output presentations could perform equally well over the whole truth table but have substantially different trained weight-matrices.

Connection weights were updated on each training cycle, after the connection-weight updates computed by the algorithm were scaled by a learning rate term. The learning rate was set to 1 for the canonical and structure weights, and to 0.10 for the non-structure weights, in order to disadvantage the non-structure connections. All weights had a lower bound of 0 except for the canonical weights, which had a lower bound of 1 to ensure their participation in canonical interactions.

Networks having full connection-weight matrices were trained, and then retrained after a pruning procedure that removed about 40% of the non-structure weights on average. Pruning was undertaken both to reduce the number of non-structure weights and to improve network performance, specifically, generalizability (see Camacho et al., 2018 for details).

### 2.4 Adjusting Adaptable TSCs

Neuroadaptation has been observed in many neural systems under many different kinds of sustained perturbations and is known to occur through changes in the “activities” (expression, sensitivity, cellular/synaptic localization, etc.) of key proteins (ion channels, transmitter receptors, etc.) (Turrigiano, 2008, 1999). Neuroadaptation was simulated in the model by adjusting the connection weights representing the 13 adjustable-TSCs, which have been found experimentally to adjust under chronic stress, antidepressant, or steroid hormone administration. The neuroadaptation process is distinct from the neural network training procedure. In neuroadaptation, only a subset of the weights (specifically the 13 adjustable-TSCs) are changing, and the weights are not changing to produce agreement with the truth table but instead are adjusting to produce a more general overall restoration of the baseline activity of the monoaminergic brain regions.

Administration of a drug or hormone can cause the activities of the units corresponding to the monoaminergic brain regions to diverge from their normative, no-input baselines. We define the sum of the absolute difference of the DR, LC, and VTA units from their baselines as the network “adaptation error”, and we define “initial error” as the adaptation error before any TSC strength adjustments have occurred. A network is considered “adjusted” if 1 or more of its TSC strengths has been adjusted, and an adjusted network is considered “adapted” if its adaptation error falls 25% or more below its initial error.

In order to account for the natural variability between patients, we studied TSC-strength adjustments in 3 different representative networks, each trained from different random initial weight matrices using different random orders of input/desired-output training-pattern presentation. We found all configurations of the 13 TSC weights reachable from each of the 3 representative networks for all increments of 1.00 within bounds of 0 to |5| up to a total of 6 adjustments. If an adjustable-TSC strength was near 0 or |5|, and an increment of 1.00 would exceed a bound, the TSC was adjusted by a partial increment to reach the bound. In this way we produced 579,125 total TSC-strength configurations over the 3 networks. Possible neuroadaptive outcomes of the MSS-model were studied by analyzing this large set of adjusted configurations.

Generation of the full set of TSC-weight configurations for a total of 7 adjustments of the 13 adjustable-TSCs would have produced up to 67,977,560 configurations. We were unable to analyze such a large set of configurations due to limitations in available computer memory (see Supplemental Material S3).

### 2.5 Full-range Individual-weight Adjustment (FRIWA)

We searched the 579,125 total TSC-strength configurations over the 3 networks for configurations that made their networks “therapeutic” to chronic SSRI. Therapeutic configurations were defined as those adapted configurations that raised 5HT up to or above the 5HT therapeutic floor (>=0.70) and decreased CORT below the therapeutic CORT ceiling (<=0.70). Of the 579,125 total TSC-strength configurations, 241,174 were adapted under chronic SSRI and of those, 12,351 were therapeutic.

It is possible that some adjustable-TSCs contribute more than others to the therapeutic state. We define “therapeuticity” as the ability of an adjustable TSC to contribute to the attainment of therapeutic monoamine or cortisol levels through changes in its strength. We conducted full-range individual-weight adjustment (FRIWA) analysis to determine the therapeuticity of individual adjustable-TSCs.

For FRIWA analysis, the weight of a single adjustable-TSC is adjusted across its full range (0 to |5|) in individual weight adjustment (IWA) increments of 1.00 (or fractional increments if the weight was near a bound), while the weights for the other 12 adaptable TSCs remained frozen at their starting values, in each of the therapeutic starting configurations. FRIWA generated a set of 78 new configurations (6 IWA adjustments for each of 13 TSC weights) starting from each of the 12,351 therapeutic configurations for a total of 963,378 new configurations. All configurations that were no longer adapted after a step of IWA were excluded from further analysis, leaving 377,939 adapted TSC-strength configurations. Of those, 354,462 configurations were classified as “resistant,” because they remained therapeutic despite a step of IWA, while the remaining 23,477 configurations were classified as “sensitive” because they were no longer therapeutic following a step of IWA.

The weights of the 13 adjustable-TSCs in all of the post-IWA configurations were pooled in each of the 3 networks and separated on the basis of resistance and sensitivity, excluding the weights that were manipulated by FRIWA. The mean weight of each adjustable-TSC in each network was then computed for both the resistant and sensitive configurations and compared. Further FRIWA analysis involved computing the pairwise correlations between all TSC weights over all resistant and sensitive configurations for each network. Analogously in the correlation analysis, the TSC weights that were manipulated by FRIWA were excluded. Only correlations that were observed over all 3 representative networks would have been reported in Results.

### 2.6 Temporal-logic Model-checking

In addition to enumerating all possible TSC-strength configurations for up to 6 adjustments, we also studied neuroadaptation as a process by making the allowed TSC-strength adjustments (increments of 1.00 up or down, within bounds of 0 and |5|) in all possible sequences. To study neuroadaptation as a process, we constructed the tree of transitions between TSC-strength configurations. The root of the tree is the TSC-strength configuration before any adjustments are made. The first level of the tree contains all degree-1 adjustments, the second level contains all degree-2 adjustments, and so on for 6 degrees of adjustment (the tree root is degree-0). Thus TSC-strength configurations evolved incrementally in time.

Linear temporal-logic (LTL) can be used to express and deduce logical propositions about systems whose “states” evolve systematically in time. Here we use LTL to evaluate propositions on temporal relationships that occur during an incremental process of neuroadaptation. In our LTL analysis, states are construed as specific TSC-strength configurations, along with the properties of a network instantiated with a specific TSC-strength configuration (i.e. adaptation error, and the levels of 5HT and CORT).

We evaluated whether a specific degree of neuroadaptation (i.e. a configuration in which a specific TSC has been adjusted up 3 times (sensitized) or adjusted down 3 times (desensitized)) always leads to therapeutic configurations, or the TSC is no longer sensitized or desensitized, for all possible sequences of adjustments proceeding from that configuration, up to a total of 6 adjustments. To view the predicates and propositions used in LTL analysis, see Supplemental Material S5. If a single adjustable-TSC mediated the therapeutic state by itself, then the model-check for either of those 2 propositions for that specific, adjustable-TSC should be True.

We also used a form of LTL to examine the proportions (percentages) of state transition sequences that met therapeutic criteria. We found all sequences proceeding from any therapeutic state, serving as a reference state, along which all subsequent states were also therapeutic (see Supplemental Material S5). We placed no restriction on the degree of adjustment in any of the reference TSC-strengths, but at least one adjustment was needed to achieve a therapeutic state because the unadjusted TSC-strength configuration (transition-tree root) was not adapted by definition. We looked for sequences of all degrees starting from reference TSC-strength configurations of degree-1 or higher, and proceeding up to degree-6 (6 total adjustments from the tree root). For no therapeutic reference TSC-strength configurations were all subsequent states therapeutic. Instead, we found the proportion of sequences for which this was true for sequences of up to 5 adjustments and reported them in Results.

## 3. Results

### 3.1 Agreement between Actual and Desired Outputs

The results of network training can be viewed in Figure 2. Each plot in Figure 2 shows all of the desired and actual outputs for a single transmitter or hormone producing region, transmitter, or hormone output unit (DR, LC, VTA, PVN, Testes, 5HT, NE, DA, CORT, or TEST). Each actual output or desired output is plotted as a solid or dashed line, respectively.

**Figure 2:**
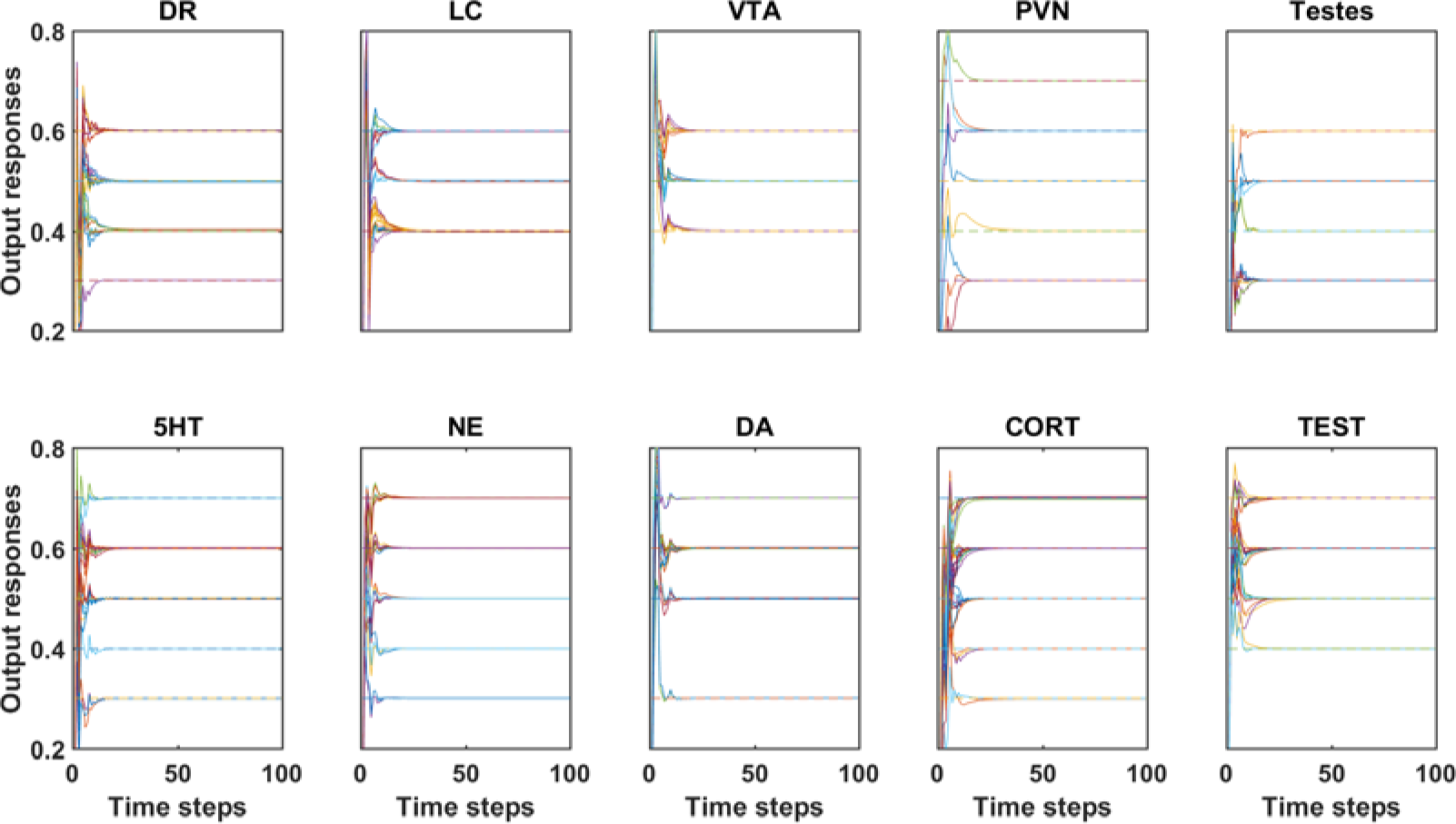
Agreement between actual model outputs and desired outputs after pruning and retraining. Single plots compare the desired (i.e., target) outputs with the actual responses of model output units representing monoamine or hormone producing regions, or the monoamines or hormones themselves (DR, LC, VTA, PVN, Testes, 5HT, NE, DA, CORT, and TEST). Actual or desired outputs are plotted as solid or dashed lines, respectively. Note that the solid lines, representing actual outputs, reach steady-states that overlie the dashed lines, representing corresponding desired outputs, illustrating the accuracy of the training procedure.

Following an initial transient of 25 time steps or less, the activity level of each unit settles into a stable activity pattern that is maintained for the duration of the run (100 time-steps). Agreement between steady-state actual and desired-output responses is nearly exact. The root-mean-square (RMS) error of this network over the entire truth-table was very low (1.84 × 10^−4^). Supplemental Figure 2 illustrates model agreement for all output units over all input/desired-output patterns for 1 representative network.

### 3.2 Enumeration of Adjustable TSC-strength Configurations

The baseline, or normative, activity level of each unit in a non-adjusted network is its response when network input is 0 (no drug, hormone, lesion, or other input). Figure 3 compares baseline activity levels (dashed blue lines) of the units representing the 3 monoaminergic regions (DR, LC, and VTA), and the units representing 5HT and CORT, with their responses to acute SSRI administration (orange lines). Deviation in the activity levels of the units from their baselines with acute SSRI produces an imbalance, or adaptation error, which can be reduced through neuroadaptation. An “adapted network” is any network instantiated with a TSC-strength configuration that is adapted to chronic drug/hormone administration (i.e. the continued presence of a drug or a drug/hormone combination). The DR, LC, VTA, 5HT, and CORT responses of an example network adapted to chronic SSRI is shown with yellow lines in Figure 3. In this example adapted network, the DR and VTA responses move back toward normative baselines while the 5HT level increases and the CORT level decreases. This pattern of elevated 5HT and reduced CORT levels has been observed in responders to chronic SSRI administration (Ceglia et al., 2004; Ruhé et al., 2015)

**Figure 3:**
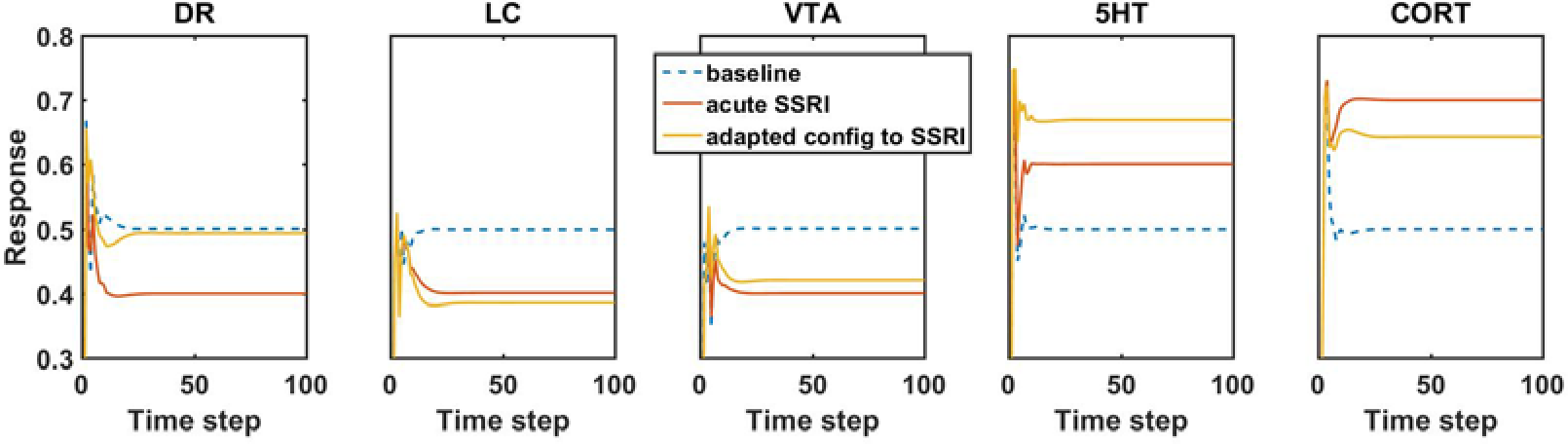
Comparison of model unit responses at baseline (no-drug), with acute (no-adaptation) SSRI administration, and with chronic (adaptation) SSRI administration. The responses of the 3 key monoaminergic brain regions (DR, LC, and VTA), as well as 5HT and CORT, are shown in the subplots as labeled. In each subplot, the blue dashed line represents the baseline (no-drug) activity level of each unit. The red line in each subplot represents the responses of the units with acute (no-adaptation) SSRI administration. Note that the responses of the 3 key brain regions all decrease and the levels of 5HT and CORT both increase in the acute SSRI condition. The yellow line in each plot shows the adapted activity levels of each unit in an example adapted configuration with chronic SSRI administration. In this adapted configuration, the DR and VTA responses return closer to baseline, and 5HT and CORT responses increase and decrease, respectively.

Not all adapted networks, however, produced this therapeutic pattern of 5HT and CORT release. To determine the possible effects of chronic administration of drug/hormone combinations on a patient population, the responses to that combination over the total of 579,125 adjusted networks were determined (note that each of the 579,125 adjusted networks has 1 of the 579,125 adjusted TSC-strength configurations). The distributions of monoamine and CORT levels over all of the networks that are adapted to specific drug/hormone combinations are shown as histograms Figures 4 and 5.

**Figure 4:**
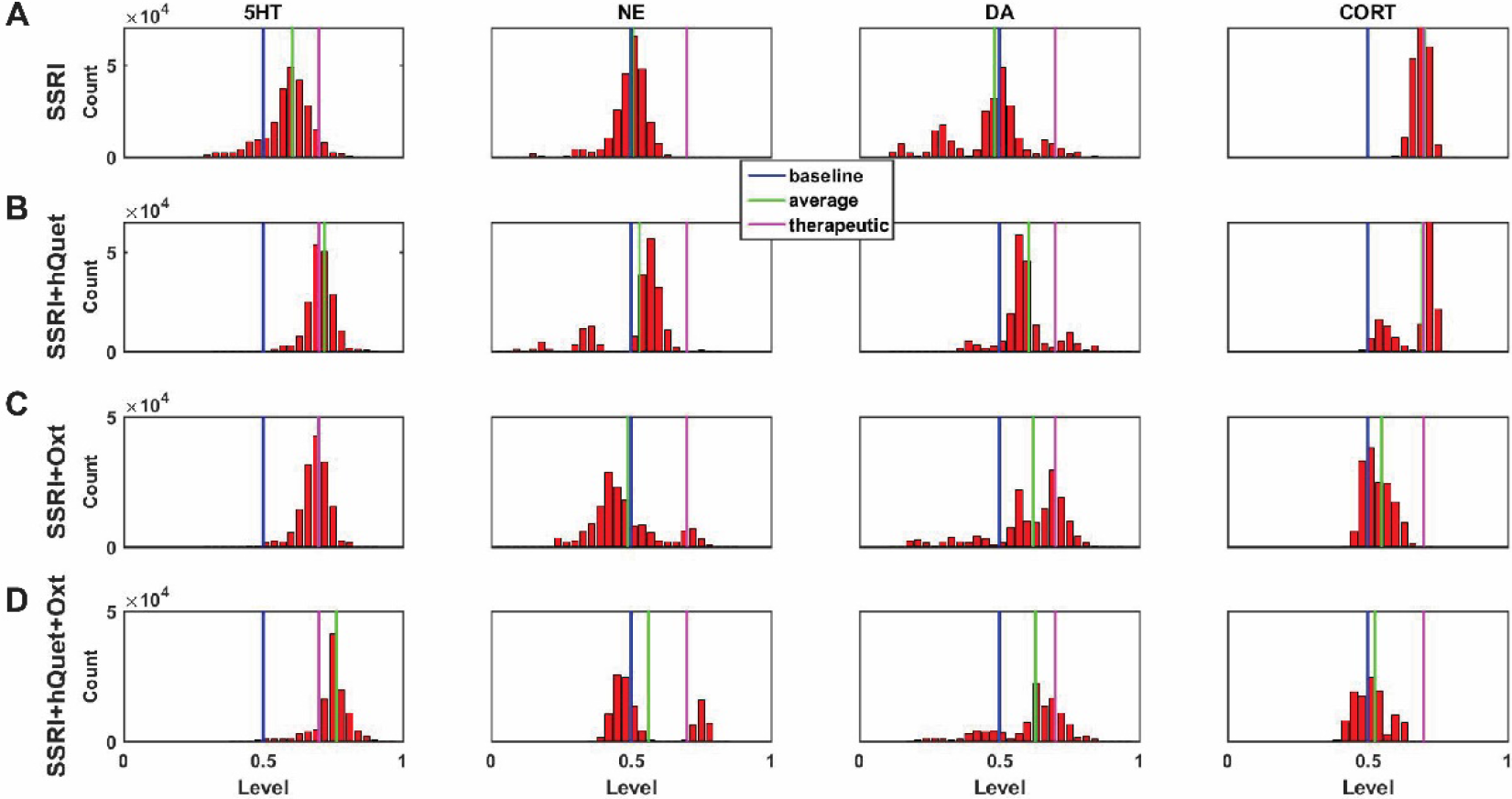
Histograms showing number of adapted configurations associated with different monoamine and cortisol levels with combinations of SSRI, Quetiapine, and Oxytocin. Networks were adapted to SSRI alone (A), SSRI+Quetiapine (hQuet) (B), SSRI+Oxytocin (Oxt) (C), and SSRI+hQuet+Oxt (D). (B) and (C) show that combining an SSRI with either hQuet or Oxt increases the proportion of high monoamine and low CORT states over the SSRI alone. (D) shows that combining an SSRI with both hQuet and Oxt further increases the proportion of high monoamine and low CORT states.

**Figure 5:**
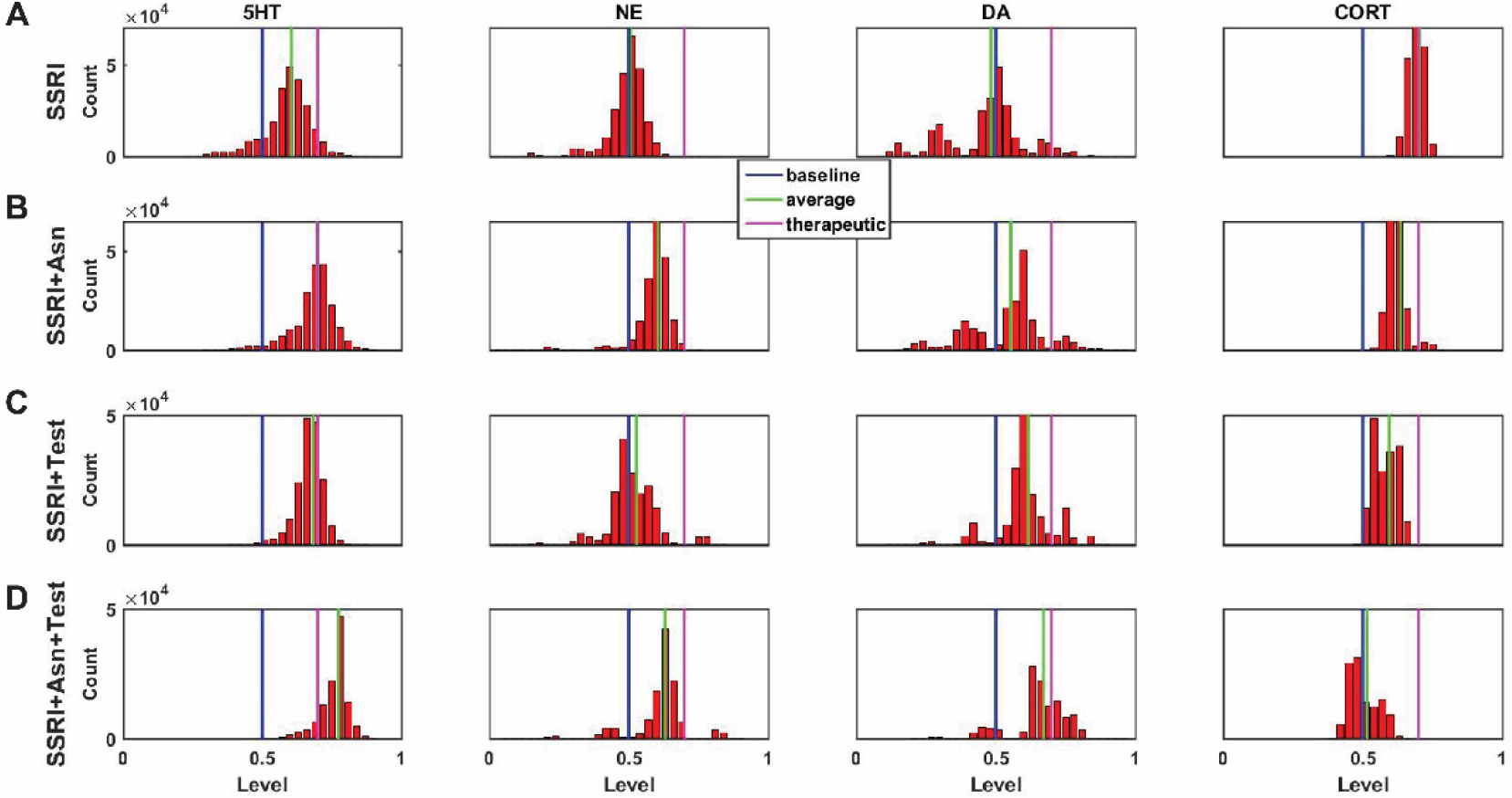
Histograms showing number of adapted configurations associated with different monoamine and cortisol levels with combinations of SSRI, Asenapine, and Testosterone. Networks were adapted to SSRI alone (A), SSRI+Asenapine (Asn) (B), SSRI+Testosterone (TEST) (C), and SSRI+Asn+TEST (D). (B) and (C) show that combining an SSRI with either Asn or TEST increases the proportion of high monoamine and low CORT states over the SSRI administered alone. (D) shows that combining an SSRI with both Asn and TEST further increases the proportion of high monoamine and low CORT states.

Each histogram in Figures 4 and 5 shows the numbers of adjusted networks (each with its associated, adjusted TSC-strength configurations) adapted to each drug/hormone combination that had levels of 5HT, NE, DA, and CORT falling within discrete bins as indicated. In all histograms the baseline, average, and therapeutic levels of each neurotransmitter or hormone is represented with a blue, green, or magenta line, respectively. Although “therapeutic” monoamine levels have not been quantified, we set the therapeutic 5HT, NE, and DA floors and the therapeutic CORT ceiling to 0.70 using estimates based on experimental findings described in Supplemental Material S4.

Comparison of Figure 4A (SSRI alone) with 4B shows that combining SSRI with Quetiapine (an antipsychotic drug) not only increases the proportion of adapted states with therapeutically elevated (toward the right) 5HT, NE, and DA, but also increases the proportion of adapted states with therapeutically decreased (toward the left) CORT. Figure 4C shows that combining SSRI with Oxytocin (a hormone) increases the proportion of adapted states with therapeutically elevated 5HT and DA as well as the proportion of states with low CORT. Figure 4D shows that the combination of all 3 factors (SSRI, Quetiapine, and Oxytocin) further increases these proportions. Figure 4D shows an interesting bi-modal distribution of NE, where one peak is at or below baseline NE levels while the second peak is shifted to the right above the NE therapeutic floor.

Comparison of Figure 5A and Figure 5B shows that combining SSRI with Asenapine (an antipsychotic drug) increases the proportion of adapted states with therapeutically elevated levels of all 3 monoamines and low CORT. Figure 5C shows that combining SSRI with TEST increases the proportion of adapted states with therapeutically high 5HT and DA and low CORT. Figure 5D shows that the combination of SSRI, Asenapine, and TEST shifts all 4 of these distributions further toward their respective therapeutic poles. These histograms illustrate how the model can be used to identify drug or drug/hormone combinations that could potentially be therapeutic for a higher proportion of male patients than SSRIs by themselves (see Discussion).

### 3.3 Predicting the Chronic Monoamine Levels Associated with Drug and Hormone Combinations

Different depressive subtypes are believed to respond better to elevations in different monoamines (Malhi et al., 2005; Parker et al., 1992). We were therefore interested in the chronic effects of different drug/hormone combinations on adapted monoamine levels. To that end, the model was used to evaluate networks adapted to chronic administration of SSRI, SSRI paired with 24 clinically relevant drugs or hormones, triples consisting of Testosterone and each of the 24 pairs, and triples consisting of Testosterone, Oxytocin, and 1 drug from each pair for a total of 74 drug/hormone combinations.

The average levels of each of the 3 monoamines over all networks adapted to each drug/hormone combination were assembled into an average monoamine vector [5HT_avg_ NE_avg_ DA_avg_]. The average monoamine vector for each drug/hormone combination is shown as a row in the heatmap in Figure 6. The following monoamine reference vectors were defined: baseline [0.50 0.50 0.50], therapeutic [0.70 0.70 0.70], and excess [0.80 0.80 0.80]. The excess monoamine reference vector is included to aid in identification of drug/hormone combinations that may elevate monoamines to toxic levels (Boyer and Shannon, 2005; Shrier et al., 2000). All adapted, average monoamine vectors [5HT_avg_ NE_avg_ DA_avg_] that elevated 1 or more monoamines above 0.80 (representing triple the baseline value) were ordered by their vector distance from the excess monoamine vector. All other adapted monoamine vectors were ordered by their vector distance from the therapeutic reference vector. This figure can help guide clinical decision-making for treatment of specific depressive subtypes that respond to elevations in specific monoamines.

**Figure 6:**
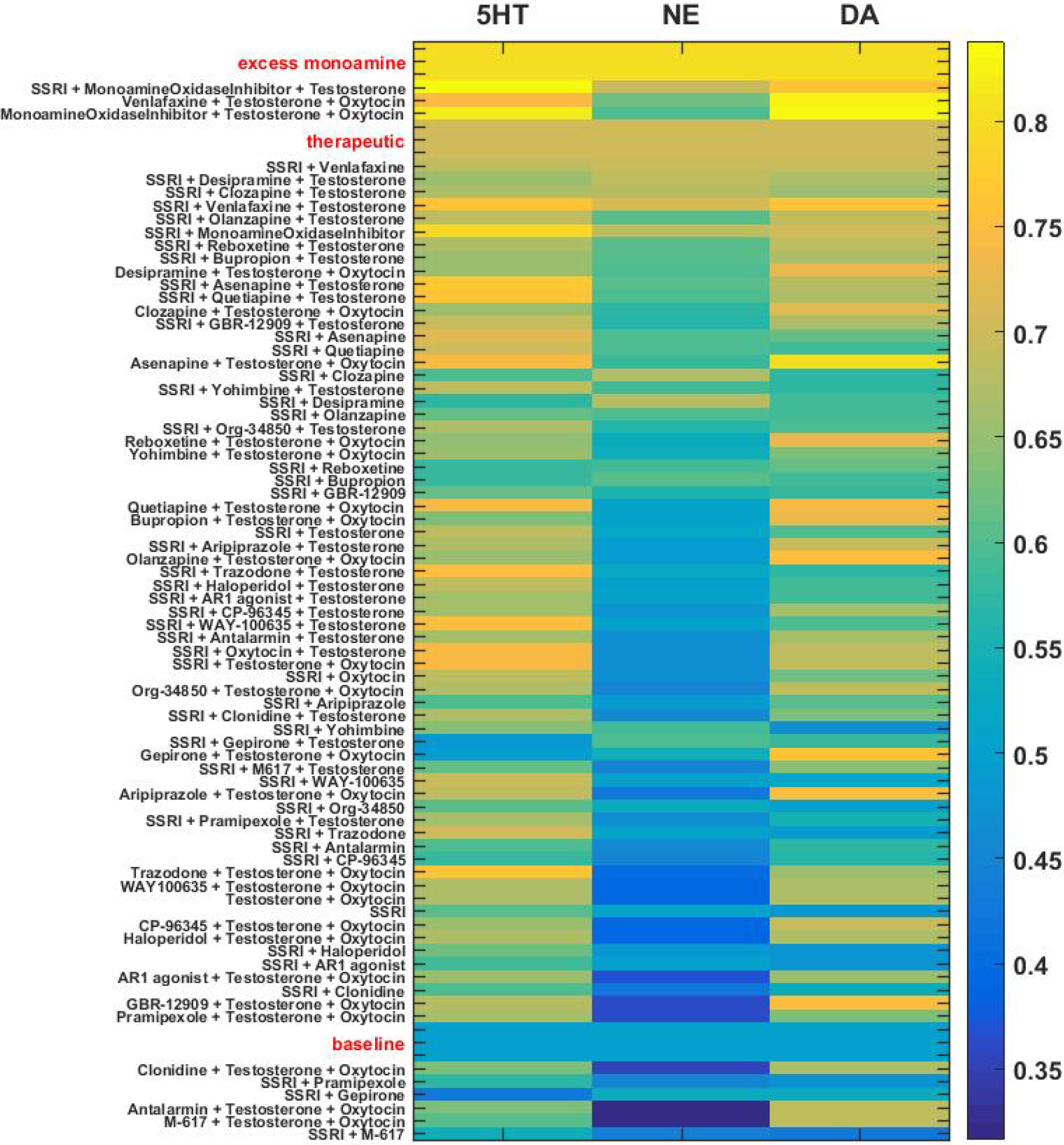
Heatmap of adapted monoamine levels with SSRI, other selected drugs or hormones paired with SSRI, and selected SSRI/other-drug/hormone triples. Adapted monoamine levels associated with neuroadaptation to SSRI, SSRI paired with other selected drugs or hormones, SSRI/other-drug/hormone triples, and drug (SSRI or other)/hormone/hormone triples were averaged over the three networks and expressed as a vector [5HT_avg_ NE_avg_ DA_avg_]. Vectors associated with drug or hormone pairs and triples, along with the baseline reference vector ([0.50 0.50 0.50]), were ordered by vector distance from the therapeutic reference vector ([0.70 0.70 0.70]). The exceptions were drug or hormone combinations that produced one or more excess monoamine levels (> 0.80), which were ordered by vector distance from the excess monoamine reference vector ([0.80 0.80 0.80]).

### 3.4 Full-range Individual-weight Adjustment (FRIWA) to Evaluate TSC Therapeuticity

In order to identify single adjustable TSCs that may mediate therapeutic effects by themselves, a FRIWA analysis was conducted that evaluated the contribution of each individual adjustable-TSC to therapeutic adaptation with chronic SSRI administration. The weights for each TSC were compiled for all resistant and all sensitive post-FRIWA configurations and averaged for each of the 3 representative networks separately.

The average values of the TSC weights over either the resistant or sensitive configurations are plotted for each representative network in blue, red, or green in Figure 7. This figure illustrates that the average resistant and sensitive TSC strengths (corresponding to TSC network connection weights) differ between the 3 representative networks, but that the average resistant and sensitive TSC strengths are about the same in each network. Although the 3 networks are indeed different in their specific weight values, the FRIWA analysis shows that they are similar in that no single TSC in any of the 3 is determinative of the therapeutic state by itself.

**Figure 7:**
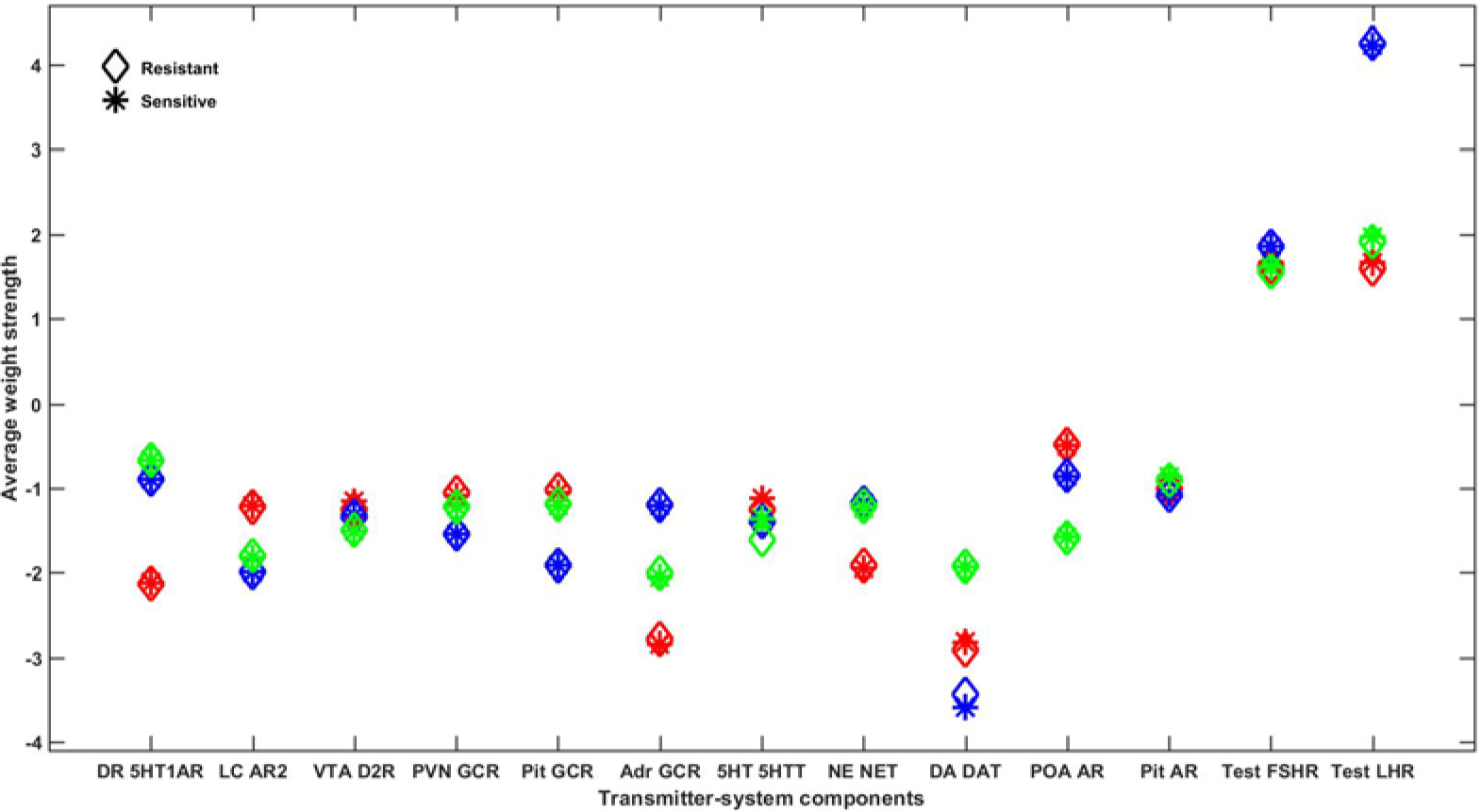
Average adjustable TSC-strength comparison between resistant and sensitive configurations. Every therapeutic configuration adapted to chronic SSRI was assessed for resistance to full-range adjustments of the weights representing each of the 13 adjustable-TSCs taken individually. The average strength of each of the 13 adjustable-TSCs in all of the configurations, excluding those in which that TSC itself was adjusted using FRIWA analysis, was computed for both the resistant and sensitive configurations and plotted as asterisks or diamonds, respectively. The results for all 3 representative networks are shown on this single plot with 3 different colors (red, blue, or green) to distinguish between the mean adjustable-TSC strengths of each network. Note that the average resistant and sensitive strengths for each adjustable-TSC are very close in all 3 networks, illustrating that no individual TSC mediates the therapeutic state by itself.

### 3.5 Pairwise Correlations between Adjustable TSC Strengths

To evaluate the possibility that the therapeutic state may be determined by correlations between pairs of TSCs rather than by individual TSCs alone, we conducted a pairwise correlation analysis over all resistant and all sensitive configurations, separately in each of the 3 representative networks. Despite the permissive statistical significance level of p = 0.05, the analysis found no significant pairwise correlations between adjustable TSCs in either the resistant or sensitive configurations that were consistent over all 3 representative networks. This result suggests that there are likely no pairs of adjustable TSCs that can pharmacologically be targeted to relieve depressive symptoms in all patients.

### 3.6 Temporal-logic Model-checking

We analyzed neuroadaptation to chronic SSRI as an incremental process using two forms of LTL analysis. We used the LEADS TO operator to determine if specific degrees of neuroadaptation in individual TSCs precede therapeutic configurations. We used the NEXT operator to determine if therapeutic configurations tend to arise from specific TSC-strength configurations.

The antecedent of the LEADS TO LTL propositions we analyzed specified that a specific TSC had been adjusted 3 times (halfway), either up (sensitized) or down (desensitized). The consequent queried whether all subsequent adjustments (after a specific TSC had been sensitized or desensitized 3 times as specified in the antecedent) in any subset of the 13 TSCs would all LEAD TO a therapeutic configuration, or the TSC loses its degree of sensitization or desensitization. All 26 LTL propositions (see Table 2) returned false in all 3 networks out to 6 total adjustments. This analysis shows that 3 adjustments of any individual adjustable TSC weight up or down does not necessarily lead to therapeutic states.

**Table 2:**
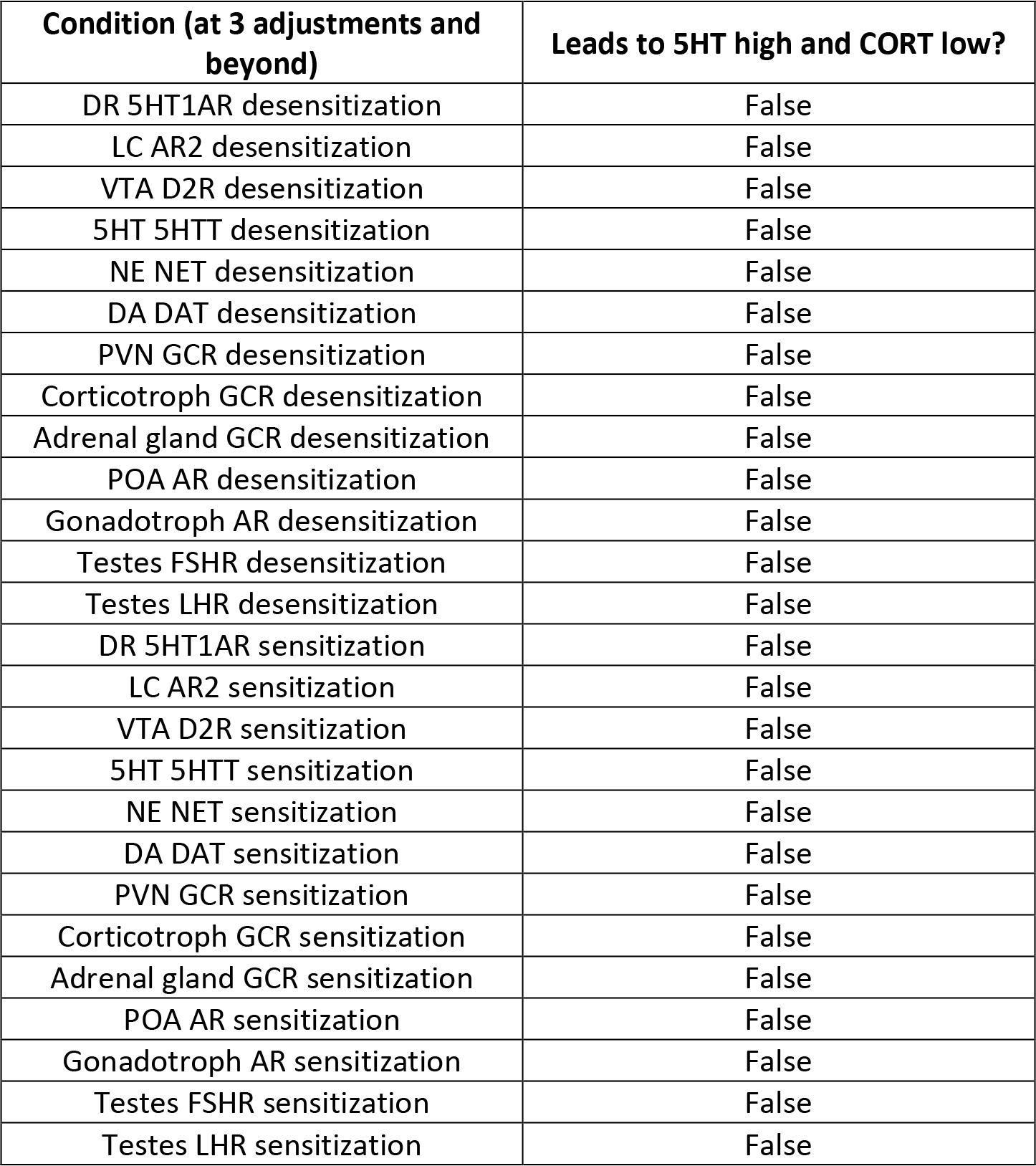
Linear temporal logic (LTL) analysis on the association between TSC-strength adjustments and the therapeutic state.

We were next interested in whether there are full or partial subtrees of adapted and therapeutic states within the tree of allowed transitions between TSC-strength configurations. We used the NEXT LTL operator to find sequences of therapeutic states. This analysis revealed that 37.80% of therapeutic configurations have the property that over 50% of subsequent TSC-strength configurations within 4 adjustment steps are also therapeutic. This analysis shows that therapeutic states tend to be spatially localized in a few subtrees within the tree of possible TSC-strength configurations.

## 4. Discussion

Clinicians designing antidepressant augmentation strategies are challenged by the overwhelming complexity of depression neurobiology, and the potentially even more overwhelming number of possible drug combinations to choose from to treat depression. Our model addresses both challenges. By incorporating known neurobiological interactions in its structure and experimental findings in its training set, the MSS-model incorporates the complexity of the monoaminergic-neurotransmitter systems, the stress-steroid system, and the male sex-steroid system in its analysis. In that it agrees in structure and function with a broad range of empirical findings, the model is a valid representation of many of the neuroendocrine interactions that underlie male depression. As such, the MSS-model allows for computational screening and analysis of a very large number of drug or drug and hormone combinations as an aid in the identification of combinations that hold promise for the clinical management of depression in men. An example of this application is computational identification of drug/hormone combinations that could be effective for patients with different subtypes of depression (Figure 6), which respond to elevations in the levels of different monoamines, by computationally identifying the monoamine profiles associated with each combination (Malhi et al., 2002; Morris et al., 2012).

Like its predecessors, the MSS-model incorporates many TSCs that are known to adjust their strengths under conditions such as chronic drug administration. In consequence of its many adjustable TSCs, and again like its predecessors, the MSS-model has many ways to neuroadapt but not all of those ways are therapeutic. Together the models suggest that real patients likewise have many ways to neuroadapt to chronic antidepressant treatment and indicate that we, therefore, should not expect them all to respond in the same way. Heterogeneity among patients in their neuroadaptive pathways may explain the relatively low efficacy rates observed in antidepressant clinical trials (Cipriani et al., 2016, 2008; Trivedi et al., 2006b; Young et al., 2009).

The MSS-model analysis included a novel LTL NEXT analysis designed to search for specific subtrees of adapted and therapeutic states within the tree of transitions between TSC-strength configurations. This analysis found that there were no therapeutic configurations, serving as reference configurations, from which all degree-1 transitions led to therapeutic configurations. Clearly then, there would be no reference TSC-strength configurations from which transitions of degree-2 or higher all lead to therapeutic configurations. However, there were reference TSC-strength configurations from which the majority of subsequent configurations were therapeutic, implying that the TSC-strength configuration space has some regions that are more therapeutic than others.

The analyses of the MSS-model and of its predecessors together characterize the process of neuroadaptation to chronic antidepressant as highly multifactorial. The level of complexity it reveals in the neurobiology of depression translates into a formidable clinical challenge in designing effective pharmacological interventions for all depressed patients. But the model also provides a means to meet that same challenge. By incorporating much of the known structure and function of the complex neuroendocrine interactions that underlie depression, the model provides a mechanism for computationally evaluating potentially effective drug or drug/hormone combinations.

This version of the MSS-model is specific to the male neuroendocrine system. It shows that certain combinations of antidepressants and the hormone testosterone could be more effective than SSRI by itself as a treatment for depression in men. Preliminary clinical studies suggest that Testosterone supplementation can enhance the antidepressant response (Kanayama et al., 2007; Miclea et al., 2018; Orengo et al., 2005). The combinations that included an SSRI, another drug that targeted the monoamines, and Testosterone were in the upper range of the Figure 6 heatmap close to the therapeutic reference vector.

The majority of the antidepressant combinations in Figure 6 show a reduction of NE levels (heatmap colors in the “blue” range) with chronic administration. It is unclear if the configurations adapted to these combinations all reduce NE, however. The histogram in Figure 4D illustrates that neuroadaptation to chronic antidepressant combinations (specifically SSRI/Quetiapine/Oxytocin in this figure) can lead to a bi-modal distribution of NE, wherein about half of the neuroadapted configurations result in reduced NE levels, and the remaining half of neuroadapted configurations result in elevated NE levels. The results in this histogram suggest that different depressed male patients may respond to chronic antidepressant administration with different patterns of NE release.

Unfortunately, data on the female neuroendocrine system sufficient to develop a female version of the MSS-model analogous to the male version was unavailable. Although the depression literature has a reasonable volume of experimental data on the female sex-steroid system and depression, scientific reports of findings on the interactions between the monoaminergic-neurotransmitter, stress-hormone, and sex-hormone systems of females are rare. This lack of data from female animal models of depression is especially problematic because women are twice as likely as men to suffer from depression (Frerichs et al., 1981; Nolen-Hoeksema, 1987; Wade et al., 2002).

Current clinical approaches to antidepressant pharmacotherapy are largely similar for both sexes (Koenig and Thase, 2009), yet it has been observed that women and men respond better to different antidepressant drugs and combinations (Khan et al., 2005; Kornstein et al., 2000). Sex-specific computational models would provide the capability to individualize antidepressant treatment regimens by computationally screening for drug and drug/hormone combinations in models that incorporate sex differences in their structure and behavior. It is therefore crucial for experimental neurobiologists to collect data on female monoaminergic-hormonal interactions, in order to permit modelization of the female neuroendocrine system and computational identification of novel drug/hormone combinations that potentially could treat depressed women more effectively.

## Contributors

MBC performed all literature searches, wrote the MATLAB computer programs, performed computer simulations and analyses, collected data, and wrote the manuscript. WDV developed LTL procedures, performed LTL analyses, and co-wrote the manuscript. TJA designed the study, developed the network modeling and machine-learning applications, wrote the initial computer programs, directed the research, and co-wrote the manuscript.

## Acknowledgments

We thank Ignacia Caviedes, Haven Comeaux, Kate Hamblen, Emily Hamm, Neena Joshi, Mahak Lalani, Diana Masolak, Cassandra Mora, and Katherine Zitello for their help in compiling the database of experimental facts on which the MS-model is based. We thank Allegra Domel for her contribution to the computational pruning analysis. We also thank Shivali Patel for her computational contribution to the construction of the full model-diagram.

## Supplementary material

Supplementary data associated with this article can be found on the publisher’s website.

