## Supplementary Material for "Computational Analysis of Therapeutic Neuroadaptation to Chronic Antidepressant in a Model of the Monoaminergic Neurotransmitter, Stress Hormone, and Male Sex Hormone Systems"

### *Supplemental Material*

**Mariam Bonyadi Camacho, Warut D. Vijitbenjaronk, Thomas J. Anastasio***

#### Supplemental Material:

This document serves as a supplement to the manuscript, **Computational Analysis of Therapeutic Neuroadaptation to Chronic Antidepressant in a Model of the Monoaminergic Neurotransmitter, Stress Hormone, and Male Sex Hormone Systems.**

#### Contents

***Text***

Abbreviations Chart 1

**S1**: Details on Model Structure 2

**S2**: Truth-table Justification 4

**S3**: Hardware Considerations 22

**S4**: Setting Therapeutic Criteria 22

**S5**: Details on LTL Analysis 22

***Figures***

**Supplemental Figure 1**: Complete model structure schematic 24

**Supplemental Figure 2**: Close agreement between desired (i.e. target) and actual output responses after training but before pruning (A−C) and after pruning and re-training (D−F) 25

***Tables***

**Supplemental Table 1**: Complete MSS-model truth-table 26

**Supplemental Table 2**: Canonical MSS-model weights 27

**Supplemental Table 3**: Adjustable MSS-model TSCs 29

***References*** 31

**Supplemental Abbreviations Chart**

| Term | Abbrevation |
|---|---|
| Adrenal gland | AG |
| Adrenergic receptor-2 | AR2 |
| Androgen receptor | AR |
| Asenapine | Asn |
| Corticotropin-releasing factor | CRF |
| Cortisol | CORT |
| Dopamine | DA |
| Dopamine beta-hydroxylase | DBH |
| Dopamine transporter | DAT |
| Dopamine-2 receptor | D2R |
| Dorsal raphe nucleus | DR |
| Estrogen | E |
| Estrogen receptor | ER |
| Estrogen receptor alpha | ER-alpha |
| Estrogen receptor beta | ER-beta |
| Follicle stimulating hormone | FSH |
| Follicle stimulating hormone receptor | FSHR |
| Full-range individual weight adjustment | FRIWA |
| gamma-Aminobutyric acid | GABA |
| Glucocorticoid receptor | GCR |
| Gonadotropin releasing hormone | GnRH |
| Gonadotropin releasing hormone receptor | GnRHR |
| Hypothalamic-pituitary-adrenal | HPA |
| Hypothalamic-pituitary-gonadal | HPG |
| Individual weight adjustment | IWA |
| Leutinizing hormone | LH |
| Leutinizing hormone receptor | LHR |
| Linear temporal-logic | LTL |
| Locus coeruleus | LC |
| Messenger ribonucleic acid | mRNA |
| Monoamine | M |
| Monoamine oxidase inhibitor | MAOI |
| Monoamine-stress | MS |
| Monoamine-stress-sex | MSS |
| Norepinephrine | NE |
| Norepinephrine transporter | NET |
| Oxytocin | Oxt |
| Paraventricular nucleus of the hypothalamus | PVN |
| Preoptic area | POA |
| Progesterone | P |

| | |
|---|---|
| Quetiapine | hQuet |
| Selective serotonin reuptake inhibitor | SSRI |
| Serotonin | 5HT |
| Serotonin receptor | 5HTR |
| Serotonin transporter | 5HTT |
| Serotonin-1A receptor | 5HT1AR |
| Serotonin-1B receptor | 5HT1BR |
| Serotonin-2A receptor | 5HT1AR |
| Testes | TS |
| Testosterone | TEST |
| Transmitter system component | TSC |
| Tryptophan hydroxylase 2 | TPH2 |
| Tyrosine hydroxylase | TH |
| Ventral tegmental area | VTA |

**S1: Details on Model Structure**

The structure of the monoamine-stress-sex model (MSS-model) is based on known interactions within and between the monoaminergic neurotransmitter systems and the stress- and sex-hormone systems. The MSS-model is an extension of our previous monoamine-stress model (MS-model). Details on monoamine-stress interactions can be found in the Supplemental Material for our MS-model article (Camacho, Vijitbenjaronk et al. 2018). Experimental findings on monoamine-stress interactions with the sex-hormone system, which were used to augment the MS-model in creating the MSS-model, are described in this section.

All excitatory, inhibitory, or unknown polarities between structure connections are represented as a +1, −1, or +2 in the structure matrix. The polarities of connections where the polarity is unknown from the literature was assigned a polarity by the training procedure.

**MSS-model Extensions to MS-model Structure**

VTA DA neurons can co-release glutamate, GABA, and corticotropin-releasing factor (CRF), so there are excitatory projections from VTA to glutamate, GABA, and CRF.

The PVN secretes GnRH, so there is an excitatory connection from the PVN to GnRH (Moore and Price 1932). GnRH binds and activates the GnRH receptor (GnRHR), so there is an excitatory connection from GnRH to GnRHR (Millar 2005). GnRHRs are excitatory on the pituitary gland, so there is an excitatory projection from GnRHR to the pituitary gland (Eidne, Sellar et al. 1992). The pituitary gland secretes the gonadotropins, luteinizing hormone (LH) and follicle-stimulating hormone (FSH) (Moore and Price 1932, Harris 1964). LH and FSH then bind to LH and FSH receptors on the gonads, so there are excitatory connections from LH to LH receptors (LHR) and FSH to FSH receptors (FSHR) (Simoni, Gromoll et al. 1997, Dufau 1998). LHR and FSHR activate the gonads, so there is an excitatory projection from LHR and FSHR to the testes (Whitelaw,

Smyth et al. 1992, Simoni, Gromoll et al. 1997, Gunnarsson, Nordberg et al. 2003). The testes produce testosterone (TEST) (de Kretser, Hedger et al. 2002). There is an excitatory projection from the testes to testosterone.

ER-beta receptors increase TPH2 mRNA expression in rodents, so there is an excitatory projection from ER-beta to TPH2 (Donner and Handa 2009). Progesterone has also been found to increase TPH2 expression, so there is an excitatory connection from progesterone receptors to TPH (Bethea, Mirkes et al. 2000). Both estrogen and progesterone have been found to decrease MAO activity in Rhesus monkeys, so there are inhibitory projections from estrogen and progesterone receptors to MAO (Gundlah, Lu et al. 2002). Both estrogen and progesterone have also been found to decrease 5HT1A gene expression in Rhesus monkeys, so there are inhibitory projections from estrogen and progesterone receptors (ER and PR, respectively) to the 5HT1A receptor (Pecins-Thompson and Bethea 1999). Estrogen receptors exist in alpha and beta forms, described here as ER-alpha and ER-beta, respectively. A study using rat models found that estrogen decreases 5HT1B receptor mRNA, so there is an inhibitory projection from estrogen receptors to 5HT1B receptors (Hiroi and Neumaier 2009). Estrogen administration in rats has been found to increase 5HT2A receptor mRNA, so there is an excitatory projection from estrogen receptors to 5HT2A receptors (Summer and Fink 1995, Sumner and Fink 1998).

Estrogen has been found to increase levels of tyrosine hydroxylase (TH) and dopamine beta-hydroxylase (DBH) mRNA through an unspecified estrogen receptor, so there are excitatory projections from both estrogen receptors to TH and DBH (Serova, Rivkin et al. 2002).
Testosterone administration to male rats has been found to increase 5HT2A receptor mRNA, so there is an excitatory projection from testosterone to 5HT2A receptors (Sumner and Fink 1998). Estrogen, progesterone, and testosterone receptors have been found in DR neurons and influence DR neuron firing with unknown polarity, so there are projections of unknown polarity from ER-alpha, ER-beta, PR, and AR to the DR (Alves, Weiland et al. 1998, Robichaud and Debonnel 2005).

Both ER-alpha and ER-beta receptors have been found to mediate the inhibitory effect of estrogen on hypothalamic neurons, so there are inhibitory projections from ER-alpha and ER-beta to the hypothalamus (Lagrange, Ronnekleiv et al. 1995, Roy, Angelini et al. 1999, Skynner, Sim et al. 1999, Herbison and Pape 2001). The inhibitory effect of progesterone on hypothalamic neurons are mediated by progesterone receptors, so there is an inhibitory projection from PR to the hypothalamus (Bashour and Wray 2012). ER-alpha mediates negative feedback on the HPG axis at the level of the hypothalamus, which is the receptor that is included as an inhibitory weight in the POA (Glidewell-Kenney, Hurley et al. 2007). The inhibitory effect of testosterone on hypothalamic neurons is mediated by androgen receptors, so there is an inhibitory projection from AR to the hypothalamus (Belsham, Evangelou et al. 1998).

Negative feedback on the HPG axis also occurs at the level of the Pituitary gland in both males by androgen and ER-alpha receptors (Scully, Gleiberman et al. 1997, Thorner, Vance et al. 1998, Cheong, Porteous et al. 2014).

Androgen receptors, progesterone receptors, ER-alpha and ER-beta have all been found on the testes (Due, Dieckmann et al. 1989, Vornberger, Prins et al. 1994, Brandenberger, Tee et al. 1998, Saunders, Fisher et al. 1998, Weil, Vendola et al. 1998, Makinen, Makela et al. 2001, Juengel, Heath et al. 2006). Testosterone provides negative feedback directly on the testes through activation of androgen receptors, so there is an inhibitory connection from the testosterone receptor to the testes (Darney and Ewing 1981). The polarity of the interaction between the estrogen and progesterone receptors on the gonads have not been determined so there are connections of unknown polarity between these receptors and the testes.

**S2: Truth-table Justification**

The MSS-model includes the truth-table input/desired-output relationships derived from male organisms in the MS-model. Experimental findings supporting all additional input/desired-output relationships are described here. The scale used to describe truth-table desired-outputs (where 0.30, 0.40, 0.50, 0.60, and 0.70 represent maximal decrease, moderate decrease, baseline, moderate increase, and maximal increase) is the same in both studies (Camacho, Vijitbenjaronk et al. 2018). The only exception in the MSS-model is that he baseline for testosterone, estrogen, and progesterone was set to 0.60, 0.40, and 0.40, respectively, to reflect the difference in the levels of these hormones in males.

<u>**SSRI**</u>

Binding of the SSRI to 5HTT leads to a doubling of extracellular 5HT in male rodents, so the target-output value for 5HT with acute SSRI was set to 0.60 (Invernizzi, Belli et al. 1992, Koch, Perry et al. 2002, Calcagno, Guzzetti et al. 2009). Because acute administration of SSRIs other than fluoxetine does not significantly change NE and DA levels in male rodents, the truth table values for these neurotransmitters were set to 0.50 (Bymaster, Zhang et al. 2002, Koch, Perry et al. 2002).

The increase in extracellular 5HT associated with acute SSRI administration has been associated with increased binding to the 5HT1A autoreceptor on DR neurons, decreasing the firing activity of these neurons in male rodents (Calcagno, Guzzetti et al. 2009). The Blier group and others have found that the DR neuron firing rate decreases by about 65% in male rodents with acute SSRI, so the truth-table value for this output was set to 0.40 (de Montigny, Chaput et al. 1990, Czachura and Rasmussen 2000, Chernoloz, El Mansari et al. 2012). The Blier group found that acute SSRI administration decreases the firing rate of LC neurons by 45% and decreases the firing rate of VTA neurons by 41% in male rodents, so the truth table values for the LC and VTA firing rates with acute SSRI were also set to 0.40 (Chernoloz, El Mansari et al. 2009, Dremencov, El Mansari et al. 2009, Chernoloz, El Mansari et al. 2012). fMRI studies indicate that amygdala activity decreases moderately in both males and females, while cortical activity does not change with acute SSRI administration in males (Mayberg, Brannan et al. 2000, Kennedy, Evans et al. 2001, Takahashi, Yahata et al. 2005, Murphy, Norbury et al. 2009). The Murphy lab found that amygdala activity decreases moderately in response to neutral faces with acute SSRI in both male and female subjects, and the Mayberg lab found that PFC activity moderately increases after 6-weeks of SSRI treatment but not after acute (1-week) SSRI treatment in male subjects. The truth table value for the amygdala with acute SSRI were set to 0.40 and the truth table value for PFC was set to 0.50.

Acute SSRI administration has been found to stimulate the rodent HPA axis by increasing CRF, ACTH, and cortisol levels while also increasing PVN activity in male rodents (Jensen, Jessop et al. 1999, Wieczorek, Schulz et al. 2001, Hesketh, Jessop et al. 2005). Specifically, 30 minutes of subcutaneous cannula citalopram administration moderately increases cortisol and ACTH levels (Jensen, Jessop et al. 1999). This lab also found that PVN activity increases moderately as measured by the increase in c-fos immunoreactive cells in the PVN with acute SSRI. The same lab did not find a significant difference in CRF mRNA with acute citalopram treatment; however, another lab found that acute citalopram moderately increases CRF levels in male rodents (Moncek, Duncko et al. 2003). The Moncek lab also found that acute citalopram produces moderate increases in ACTH levels and very large increases in cortisol levels in males. The truth table values for ACTH and cortisol with acute SSRI was set to 0.60 and 0.70, respectively, in the MSS-model. The truth table value for CRF with acute SSRI was set to 0.60. Oxytocin levels have been found to moderately increase with acute SSRI, while arginine-vasopressin (AVP) levels have been found to stay the same in male rodents (Hesketh, Jessop et al. 2005). We set the oxytocin target to 0.60 and the AVP target to 0.50 with acute SSRI for males. Acute fluoxetine injection moderately increases galanin mRNA levels in male rodents, so the truth-table value for galanin with acute SSRI was set to 0.60 for males (Kuteeva, Wardi et al. 2008). Acute SSRI administration in male humans had no effect on testosterone levels, so the truth-table value for testosterone with acute SSRI for males was set to 0.60 (Schlosser, Wetzel et al. 2000).

**<u>Nomifensine</u>**

The Blier group found that acute administration of Nomifensine increases DR neuron activity by 50%, decreases VTA neuron activity by 39%, and decreases LC neuron activity by 71% in male rodents (Katz, Guiard et al. 2010). The truth-table values for the acute effect of Nomifensine on DR, LC and VTA neurons were set to 0.60, 0.40, and 0.40, respectively. The Masana study also found that acute Nomifensine maximally increases DA levels in males. The Carboni lab found that acute Nomifensine maximally increases DA levels in male rodents (Carboni, Imperato et al. 1989). The Butcher lab found a moderate increase in DA levels with acute Nomifensine in male rats (Butcher, Fairbrother et al. 1988). The truth-table value for DA with acute Nomifensine was set to 0.60.

**Reboxetine**

Reboxetine is a selective NET blocker (Hajos, Fleishaker et al. 2004). The Blier group found that acute Reboxetine administration decreases LC firing rate by 68% without affecting DR firing rate in male rodents (Szabo and Blier 2001). They also found that acute Reboxetine administration decreases VTA neuron firing rate by 31% in male rodents (Katz, Guiard et al. 2010). The truth-table values for acute Reboxetine for DR, LC, and VTA were therefore set to 0.50, 0.40, and 0.40, respectively. Acute Reboxetine administration moderately increases NE and DA levels while producing no change in 5HT levels in male rodents, corresponding to values of 0.60, 0.60, and 0.50 in the truth table for these neurotransmitters, respectively (Page and Lucki 2002). One fMRI study in both male and female humans shows that acute Reboxetine administration moderately decreases amygdala response to neutral stimuli, so the truth-table value for the amygdala with acute Reboxetine was set to 0.40 (Onur, Walter et al. 2009). Acute Reboxetine administration in male humans has been found to moderately increase ACTH levels in two studies using male volunteers, so the truth-table value for ACTH with acute Reboxetine was set to 0.60. Acute Reboxetine administration has also been shown to moderately increase cortisol levels in two different studies using male human volunteers, so the truth-table value for cortisol with acute Reboxetine was set to 0.60 (Hennig, Lange et al. 2000, Schule, Baghai et al. 2004).

**Trazodone**

Trazodone blocks 5HTT and interacts with multiple receptors of the monoaminergic system (Stahl 2009, Stahl 2009). The Blier lab found that acute Trazodone administration decreases DR neuron firing by 65%, increases LC neuron firing by 25%, and does not alter VTA neuron firing in male rodents (Ghanbari, El Mansari et al. 2010, Ghanbari, El Mansari et al. 2012). We set the target output values for DR, LC, and VTA with acute Trazodone to 0.40, 0.60, and 0.50, respectively. Acute Trazodone administration has been found to moderately increase 5HT levels without changing NE levels in male rodents, so the truth-table values for these were set to 0.60 and 0.50, respectively (Rowbotham, Jones et al. 1984, Pazzagli, Giovannini et al. 1999).

**Asenapine**

Asenapine is an antipsychotic drug that interacts with multiple receptors of the monoaminergic neurotransmitter system (Franberg, Wiker et al. 2008, Ghanbari, El Mansari et al. 2009). The Blier group found that acute Asenapine administration decreases DR neuron firing by about 30% without affecting the firing rates of the LC or VTA in male rodents (Oosterhof, El Mansari et al. 2015). We set the truth table values for DR, LC, and VTA to 0.40, 0.50, and 0.50, respectively. Acute Asenapine administration in male rats has been found to moderately increase 5HT, NE, and DA levels (Franberg, Marcus et al. 2009). The truth table outputs for 5HT, NE, and DA with acute Asenapine were all set to 0.60.

**Aripiprazole**

Aripiprazole is an antipsychotic drug that interacts strongly with DA and 5HT receptors (Shapiro, Renock et al. 2003). The Blier group found that acute Aripiprazole administration increases DR neuron firing rate by 48%, without affecting the firing rates of LC or VTA neurons in male rodents (Chernoloz, El Mansari et al. 2009). The truth-table values for the DR, LC and VTA were set to 0.60, 0.50 and 0.50, respectively. Acute Aripiprazole administration has been found to moderately increase DA levels without affecting 5HT, NE, or cortisol levels in male rodents (Li, Ichikawa et al. 2004, Zocchi, Fabbri et al. 2005, Assie, Carilla-Durand et al. 2008). The truth-table values for 5HT, NE, DA, and cortisol were set to 0.50, 0.50, 0.60, and 0.50 with acute Aripiprazole, respectively.

**Bupropion**

Bupropion is an atypical antidepressant that blocks NET and DAT (Cooper, Wang et al. 1994, Stahl, Pradko et al. 2004). The Blier group found that acute Bupropion administration moderately increases DR neuron firing rate, moderately decreases LC neuron firing rate, and does not change VTA neuron firing rate in male rodents (El Mansari, Ghanbari et al. 2008). The truth-table values for the DR, LC and VTA were set to 0.60, 0.40 and 0.50, respectively, for males with acute Bupropion administration. Acute Bupropion administration has been found to moderately increase DA and NE levels without affecting 5HT levels in male rodents (Piacentini, Clinckers et al. 2003). The truth-table values for 5HT, NE, DA were set to 0.50, 0.60, and 0.60 with acute Bupropion in the MSS-model.

**Quetiapine**

Quetiapine is an antipsychotic drug with multiple receptor and transporter affinities, including the dopamine D2 receptor, the 5HT2A receptor, and the α-1 receptor (DeVane and Nemeroff 2001, Jensen, Rodriguiz et al. 2008). The Blier group found that acute Quetiapine administration decreases the DR neuron firing rate by 43% and increases the LC neuron firing rate by 40% in male rodents (Chernoloz, El Mansari et al. 2012). The truth-table values for the DR and LC with acute Quetiapine were set to 0.40 and 0.60, respectively. Another group found that acute Quetiapine moderately increases VTA firing rate in males, so the truth-table value for the VTA with acute Quetiapine was set to 0.60 for males (Werkman, Olijslagers et al. 2004). Denys et al found that acute Quetiapine moderately increases 5HT and DA levels in the PFC, while Silverstone et al found that acute Quetiapine has no effect on 5HT levels in the PFC, but moderately increases NE and DA levels (Denys, Klompmakers et al. 2004, Silverstone, Lalies et al. 2012). Both of these studies were in male rodents. Because the Denys et al study examined multiple brain regions and also found a moderate increase in 5HT levels in the dorsal striatum with acute Quetiapine, the truth table value for DA was set to 0.60. The truth table values for 5HT and NE with acute Quetiapine were also set to 0.60.

**Pramipexole**

Pramipexole (PPX) is a D2, D3, and D4 receptor agonist (Mierau, Schneider et al. 1995). The Blier group found that acute PPX administration does not change DR neuron firing rate, decreases LC neuron firing rate by 33%, and decreases VTA neuron firing rate by 40% in male rodents (Chernoloz, El Mansari et al. 2009). The truth-table values for the DR, LC and VTA were set to

0.50, 0.40 and 0.40, respectively, with acute PPX administration, for males.

**GBR-12909**

GBR-12909 (GBR) is a DA transporter blocker (TOCRIS , Andersen 1989, Singh 2000). The Blier group found that acute GBR administration does not change DR or LC neuron firing rate but decreases VTA neuron firing rate by 26% in male rodents (Katz, Guiard et al. 2010). Another group also found a moderate decrease in VTA neuron firing rate with acute GBR in male rodents (Choong and Shen 2004). The truth-table values for the DR, LC and VTA were set to 0.50, 0.50 and 0.40, respectively, with acute GBR administration. Three labs have found moderate increases in DA levels with acute GBR in male rodents, so the truth-table value of DA with acute GBR was set to 0.60 (Rothman, Mele et al. 1991, Choong and Shen 2004, Masana, Bortolozzi et al. 2011).

**Clozapine**

Clozapine is an antipsychotic drug that targets many monoaminergic receptors (Meltzer 1994). Acute administration of Clozapine has been found to moderately decrease the firing rate of DR neurons, moderately increase the firing rate of LC neurons, moderately increase the firing rate of VTA neurons, and moderately increase the firing rate of PFC neurons in male subjects (Souto, Monti et al. 1979, Sprouse, Reynolds et al. 1999, Chen and Yang 2002, Gao 2007). Another group found that acute Clozapine completely inhibits DR firing activity in male rodents; however, the dose that was used was much higher than the standard rodent dose (Gallager and Aghajanian 1976). The truth-table values for DR, LC, VTA, and PFC were set to 0.40, 0.60, 0.60, and 0.60, respectively. Acute Clozapine administration has been found to maximally increase NE levels and moderately increase DA levels in male rodents (Zocchi, Fabbri et al. 2005). Another group also found that Clozapine moderately increases DA levels in male rodents (Masana, Bortolozzi et al. 2011). One study found that Clozapine moderately increases cortisol levels in male humans, so the truth table value for NE was set to 0.70 and the truth table values for DA and cortisol were set to 0.60 (Lee, Woo et al. 2001). One group has found that Clozapine decreases 5HT in the nucleus accumbens, while another found that Clozapine increases 5HT in the nucleus accumbens and the PFC in male rodents (Ferre and Artigas 1995, Ichikawa, Kuroki et al. 1998). Because these two groups found opposing effects of Clozapine on 5HT, and a third group found that 5HT does not change in the PFC with acute Clozapine, we used a truth-table value of 0.50 for 5HT with acute Clozapine (Zocchi, Fabbri et al. 2005).

**Ketamine**

Ketamine is an NMDA receptor antagonist (Hall and Murdoch 1990). The Blier group found that acute Ketamine administration does not change DR or VTA neuron firing rate but increases LC neuron firing rate by 23% in male rodents (El Iskandrani, Oosterhof et al. 2015). The truth-table values for the DR, LC and VTA were set to 0.50, 0.60, and 0.50, respectively, with acute Ketamine administration. One group found that Ketamine moderately increases PFC neuron activity and moderately increases extracellular glutamate while another group found that Ketamine moderately increases anterior cingulate glutamate, so the truth-table value for the PFC and glutamate were set to 0.60 (Razoux, Garcia et al. 2007, Stone, Dietrich et al. 2012, Pehrson and Sanchez 2014, Bjorkholm, Franberg et al. 2015). These studies were done in male rodents and humans. One study found that Ketamine moderately increases PFC 5HT levels in male rodents, so the truth table value for 5HT was set to 0.60 with acute Ketamine for males (Nishitani, Nagayasu et al. 2014). Two groups found that Ketamine has no effect on GABA levels in male rodents, so the truth table value for GABA was set to 0.50 with acute Ketamine for males (Lindefors, Barati et al. 1997, Stone, Dietrich et al. 2012). One lab found that acute Ketamine had no effect on male rabbit whole brain NE or NE levels in any of the brain areas examined, so the truth table value for NE with acute Ketamine was set to 0.50. The same lab also found that DA levels did not change in male rabbit whole brain with Ketamine, but did find a moderate increase in thalamus and hypothalamus DA (Glisson, el-Etr et al. 1976). Another group found that acute Ketamine moderately increases DA levels in rat PFC (Lindefors, Barati et al. 1997). Because whole brain neurotransmitter level change is the standard criterion for the truth table, the truth-table value for DA with Ketamine was set to 0.50.

**Reserpine and Reserpine/Bupropion**

Reserpine depletes monoamines by inhibiting the activity of the vesicular monoamine transporter 2 (VMAT2), which traffics monoamines to the cell membrane for extracellular release (Scherman and Henry 1984, Rudnick, Steiner-Mordoch et al. 1990). Because VMAT2 is not an element in our model, Reserpine sends inhibitory projections directly to 5HT, NE and DA directly. One group found that Reserpine moderately increases DR activity, then suppresses DR activity after about 30 minutes in male rodents (Baraban, Wang et al. 1978, Baraban and Aghajanian 1980). Because we are interested in the immediate, acute effect of Reserpine, we set the truth-table value for DR with acute Reserpine to 0.60.

**Venlafaxine**

Venlafaxine is a selective serotonin-norepinephrine reuptake inhibitor (SNRI) (Roseboom and Kalin 2000). The Blier group found that Venlafaxine decreases DR firing rate by 47% and decreases LC firing rate by 21% in male rodents, so the truth-table values for DR and LC were both set to 0.40 (Gartside, Umbers et al. 1997). Because acute Venlafaxine administration moderately increases the firing activity of the PFC in a c-fos study with male rodents, we set the truth-table value for PFC with Venlafaxine to 0.60 (Higashino, Ago et al. 2014). Higashino et al found that acute Venlafaxine administration moderately increases 5HT in the PFC, moderately increases NE in the PFC, and maximally increases DA in the PFC and Beyer et al found that acute Venlafaxine has no effect on forebrain 5HT levels but maximally increases forebrain NE levels (Beyer, Boikess et al. 2002, Higashino, Ago et al. 2014). Both of these studies were in male rodents. The truth-table value for 5HT was set to 0.60 in males to reflect a moderate increase in 5HT due to the observation by Higashino et al that 5HT increases in the PFC with Venlafaxine, and the fact that Venlafaxine is a 5HTT blocker, which are known to elevate 5HT (El Mansari, Sanchez et al. 2005). The truth-table values for NE and DA were both set to 0.70 in males to reflect the maximal increases in the levels of these neurotransmitters with acute Venlafaxine observed by the

Higashino and Beyer groups.

**Desipramine**

Desipramine is a tricyclic antidepressant with strong affinity for NET and weaker affinity for the 5HTT and DAT as well as various monoaminergic receptor targets (Berti and Shore 1967). DR firing was found to moderately decrease with acute Desipramine in male rodents, so the truth-table value for DR was set to 0.40 with acute Desipramine (Gartside, Umbers et al. 1997). Acute Desipramine also moderately decreases LC neuron activity in male rodents, so the truth-table value for LC was set to 0.40 (Scuvee-Moreau and Dresse 1979). Acute administration of Desipramine was found by Beyer et al to maximally increase NE levels without changing 5HT levels, and Kreiss et al also found that Desipramine has no effect on 5HT levels (Kreiss and Lucki 1995, Beyer, Boikess et al. 2002). Both of these studies were in male rodents. Another group found no change in 5HT levels with acute Desipramine in male rodents, but found a maximal increase in NE levels and a moderate increase in DA levels (Higashino, Ago et al. 2014). The truth-table value for 5HT was set to 0.50, for NE was set to 0.70, and for DA was set to 0.60. Acute Desipramine was found to have no effect on PFC activity in male rodents, so this truth-table value was set to 0.50 (Higashino, Ago et al. 2014). Desipramine injection in both male and female humans moderately increases blood cortisol levels after 30 minutes, so the truth-table value for cortisol with Desipramine was set to 0.60 (Asnis, Halbreich et al. 1985). Desipramine moderately increases ACTH levels in males so the truth-table value for ACTH was set to 0.60 with acute Desipramine (Kuhn and Francis 1997).

**CP-96345 and CP-96345/Stress**

CP-96345 (CP) is a neurokinin-1 receptor antagonist (Fong, Yu et al. 1992). The Blier group found that acute administration of CP in male rodents increases the firing rate of DR neurons by 46% but does not affect the firing rate of LC neurons, so the truth-table values for DR and LC were set to 0.60 and 0.50 for the DR and LC with acute CP, respectively (Conley, Cumberbatch et al. 2002, Haddjeri and Blier 2008). One group found that NK1 receptor antagonists moderately increase VTA firing rate in male rodents, so the truth-table value for VTA with acute CP was set to 0.60. The same group also found that NK1 receptor antagonists have no effect on 5HT levels in male rodents, so the 5HT value with acute CP was set to 0.50 (Lejeune, Gobert et al. 2002). Another group found that acute CP has no effect on NE levels, but moderately increases DA levels in the PFC of male gerbils (Renoldi and Invernizzi 2006). DA levels moderately increase in male rodents with acute NK1 receptor antagonists (Lejeune, Gobert et al. 2002). The truth table value for NE was set to 0.50 and for DA was set to 0.60 with acute CP. The combination of NK1 antagonists and stress leads to no change in NE or DA levels in male rodents (Renoldi and Invernizzi 2006). The truth-table values for NE and DA with CP/Stress were set to 0.50.

**Gepirone, Gepirone/Stress, and Gepirone/Dexamethasone**

Gepirone is a 5HT1A receptor agonist (Blier and Ward 2003). The Blier group and others have found that acute administration of 5HT1A agonists maximally decrease DR neuron firing rate in male rodents, so the truth-table value of DR with Gepirone was set to 0.30 (VanderMaelen, Matheson et al. 1986, Blier and de Montigny 1987, Blier and de Montigny 1990). 5HT1A receptor agonists have been found to maximally decrease 5HT levels in male rodents by multiple groups, so the truth table value for 5HT with acute Gepirone was set to 0.30 (Rutter, Gundlah et al. 1994, Dawson, Nguyen et al. 2002). 5HT1A receptor agonists have also been found to moderately increase CRF, ACTH, and cortisol release in male rodents, so the truth-table values for CRF, ACTH, and cortisol with acute Gepirone were set to 0.60 (Pan and Gilbert 1992, Matheson, Knowles et al. 1997). When males are castrated, their cortisol response to 5HT1A receptor agonists reach maximal levels, so the cortisol level for castrated males with gepirone was set to 0.70 (Matsuda, Nakano et al. 1991). The combination of Gepirone and stress moderately increases cortisol levels in male rodents, so the cortisol level in the truth-table with acute Gepirone/stress was set to 0.60 (Matheson, Knowles et al. 1997). The combination of Gepirone/Dexamethasone moderately decreases cortisol levels in male rodents, so the cortisol level in the truth-table with acute Gepirone/Dexamethasone was set to 0.40 (Matheson, Knowles et al. 1997).

**Org-34850 and Org-34850/Stress**

Org34850 (Org) is a glucocorticoid receptor antagonist (Reynolds, Saunders et al. 2015). Acute Org administration by itself has been found to have no effect on cortisol, 5HT, or DA levels in male rodents, so the truth-table values for cortisol, 5HT, and DA with acute Org were set to 0.50 (Spiga, Harrison et al. 2007, Spiga, Harrison et al. 2008). Acute **Org/Stress** was also found by this group to have no effect on cortisol levels in male rodents, so the truth-table value for cortisol with acute Org/Stress was set to 0.50.

**Oxytocin and Oxytocin/Stress**

Oxytocin is an input that projects to model oxytocin receptors (Gimpl and Fahrenholz 2001). Oxytocin administration to male rats has been shown to double DA levels, so the truth-table value for DA was set to 0.60 (Melis, Melis et al. 2007). Oxytocin by itself as well as **Oxytocin/Stress** has no effect on PVN CRF mRNA in male rats, so the CRF value with Oxytocin and with Oxytocin/Stress was set to 0.50 (Bulbul, Babygirija et al. 2011). Oxytocin has been found to moderately decrease amygdala response to fearful stimuli (faces and scenery) in human males, so the truth-table value for amygdala with Oxytocin/Stress was set to 0.40 (Kirsch, Esslinger et al. 2005). One study in rats from 1984 found that Oxytocin does not have a significant effect on ACTH levels, however, because several other groups since 1984 have found that Oxytocin moderately decreases ACTH in both male and female subjects, the truth-table values for ACTH with Oxytocin were set to 0.40 (Gibbs, Vale et al. 1984, Chiodera and Coiro 1987, Parker, Buckmaster et al. 2005). It has also been found that Oxytocin moderately decreases cortisol levels in male humans, so the truth-table value for cortisol with Oxytocin was set to 0.40 (Legros, Chiodera et al. 1984).

**Dexamethasone**

Dexamethasone is a corticosteroid with significant in vivo affinity for glucocorticoid receptors but not mineralocorticoid receptors (Bamberger, Bamberger et al. 1995, Reul, Gesing et al. 2000,

Pariante and Miller 2001). One group found that Dexamethasone maximally decreases cortisol levels in males, so the truth-table value for cortisol with Dexamethasone was set to 0.30 (Heuser, Gotthardt et al. 1994, Rush, Giles et al. 1996). Dexamethasone maximally suppresses PVN CRF mRNA in male rodents, so the truth-table value for CRF with Dexamethasone was set to 0.30 (Kovacs and Mezey 1987). Acute Dexamethasone administration moderately decreases ACTH levels in male humans, so the truth-table value for ACTH with Dexamethasone was set to 0.40 (Hohnloser, Von Werder et al. 1989). Acute Dexamethasone moderately decreases AVP mRNA in the PVN of male rodents, so the truth-table value for AVP with Dexamethasone was set to 0.40 (Kovacs and Makara 1988). C-fos mRNA responses in the PVN and pituitary gland were found to be moderately decreased with acute Dexamethasone in male rodents, so the truth-table values for the PVN and pituitary gland were set to 0.40 (Karssen, Meijer et al. 2005). Dexamethasone administration has been shown to moderately increase the levels of 5HT precursors and 5HT in male rodents, so the truth-table values for Trp, 5HTP, and 5HT were set to 0.60 with Dexamethasone (Tsubota, Adachi et al. 1999, Clark, Flick et al. 2008). Dexamethasone has also been found to moderately increase the levels of extracellular DA in male rodents but moderately decrease extracellular NE in both male and female humans, so the truth-table values for DA and NE were set to 0.60 and 0.40, respectively (Stene, Panagiotis et al. 1980, Tsubota, Adachi et al. 1999). Dexamethasone administration in male rodents has been found to decrease testosterone levels, so the truth-table value for testosterone with Dexamethasone was set to 0.50 (Saez, Morera et al. 1977).

**WAY-100635 and WAY-100635/Gepirone**

WAY-100635 (WAY) is a 5HT1A receptor antagonist (Fletcher, Forster et al. 1996). Administration of WAY has been shown to reverse the inhibitory effects of 5HT1A receptor agonists on DR firing rate in both male and female rodents, so the truth-table value for DR with WAY/Gepirone was set to 0.50 (Evrard, Laporte et al. 1999). WAY/Gepirone together has also been shown to prevent the rise in ACTH observed with Gepirone in male rodents, so the truth-table value for ACTH with WAY/Gepirone was set to 0.50 (Fletcher, Forster et al. 1996). 5HT1AR antagonists by themselves moderately increase extracellular 5HT in male rodents, so the truth-table value for 5HT with WAY was set to 0.60 (Arborelius, Nomikos et al. 1996).

**MAOI**

Monoamine Oxidase Inhibitors (MAOI) block the activity of monoamine oxidase, which is an enzyme that breaks down monoamines (Stein 1960, Remick and Froese 1990). The Blier group found that acute MAOI administration moderately decreases DR and LC neuron firing without affecting VTA neuron firing in male rodents, so the truth-table values for DR and LC were set to 0.40 and the truth table value for VTA was set to 0.50 with MAOI for males (Blier and de Montigny 1985, Chenu, El Mansari et al. 2009). Acute MAOI administration has been shown to produce maximal increases in all three of the monoamines in male rodents, so the truth-table values for 5HT, NE and DA were set to 0.70 (Butcher, Fairbrother et al. 1990, Celada and Artigas 1993, Kitaichi, Inoue et al. 2006).

**M-617**

M-617 inhibits galR1 receptors (Sevcik, Finta et al. 1993, Larm, Shen et al. 2003, Wang, Li et al. 2016). Acute M617 administration moderately decreases DR and LC firing rate and moderately decreases 5HT and NE levels in male rodents (Jacobs, Wise et al. 1974, Azmitia and Segal 1978, Seutin, Verbanck et al. 1989, Sevcik, Finta et al. 1993, Yoshitake, Reenila et al. 2003, Hawes, Brunzell et al. 2005, Mazarati, Baldwin et al. 2005). We therefore set the truth-table values for DR, LC, 5HT, and NE to 0.40. It has also been found that M617 administration produces moderate increases in the activity of the amygdala and PVN as measured by increases in c-fos expression in these regions of male rodents, so the truth-table values for these regions were set to 0.60 (Blackshear, Yamamoto et al. 2007).

**CRF1R Antagonist and CRF1R antagonist/Stress**

CRF1 receptor antagonists block the CRF1 receptor (Holsboer and Ising 2008). Specifically, multiple labs have found that CRF1R antagonists moderately decrease ACTH and cortisol release in both male and female subjects, so the truth-table values for ACTH and cortisol with CRF1R antagonist were set to 0.40 (Broadbear, Winger et al. 2004, Jutkiewicz, Wood et al. 2005, Ising and Holsboer 2007). Acute administration of CRF1R antagonist/Stress moderately increases ACTH and cortisol in male rodents, so the truth-table values for ACTH and cortisol for CRF1R antagonist/stress was set to 0.60 for males (Deak, Nguyen et al. 1999, Jutkiewicz, Wood et al. 2005).

**Haloperidol and Haloperidol/Bupropion**

Haloperidol is a dopamine D2 receptor antagonist (Schotte, Janssen et al. 1993). Subcutaneous haloperidol administration in rats produces no change in PFC NE or DA levels in male rodents, so the truth table values for NE and DA with acute Haloperidol were set to 0.50 (Li, Perry et al. 1998). The combination of Haloperidol and Bupropion produces no change in the firing rate of VTA neurons in male rodents, so the truth-table value for VTA with Haloperidol/Bupropion was set to 0.50 (Cooper, Wang et al. 1994).

**Olanzapine**

Olanzapine is dopamine D2 receptor antagonist and 5HT2A receptor antagonist (Pilowsky, Busatto et al. 1996). Subcutaneous Olanzapine administration in male rats moderately increases PFC NE and DA levels, so the truth table values for NE and DA with acute Olanzapine were set to 0.60 (Li, Perry et al. 1998).

**Clonidine**

Clonidine is an α-2 receptor agonist (Unnerstall, Kopajtic et al. 1984). Administration of Clonidine has been found to moderately decrease LC neuron firing in male rodents, and moderately decrease NE levels in both male and female humans (Svensson, Bunney et al. 1975, Veith, Best et al. 1984, Jacobs 1986). The truth-table value for LC with acute Clonidine was set to 0.40. The truth-table value for NE with acute Clonidine was set to 0.40.

**Yohimbine, Yohimbine/Stress and Yohimbine/CRF**

Yohimbine is an α-2 receptor antagonist (Perry and U'Prichard 1981). Yohimbine administration in male rats moderately elevates NE levels in the amygdala, so the truth-table value for NE with acute Yohimbine was set to 0.60 (Khoshbouei, Cecchi et al. 2002). The combination of Yohimbine and Stress maximally increases NE levels in the amygdala of male rats, so the truth-table value for NE with acute Yohimbine/Stress was set to 0.70 (Khoshbouei, Cecchi et al. 2002). The combination of Yohimbine/Stress was also found to moderately elevate galanin levels in male rats, so the truth-table value for galanin with Yohimbine/Stress was set to 0.60 (Khoshbouei, Cecchi et al. 2002). The combination of an alpha-2 receptor antagonist and CRF produces moderate increases in ACTH and cortisol in both males and females, so the truth-table values for ACTH and cortisol with Yoh/CRF were set to 0.60 (Kizildere, Gluck et al. 2003).

**Noradrenergic alpha-1 receptor agonist**

Administration of an alpha-1 agonist in male subjects moderately increases AVP levels, so the truth-table value for AVP with alpha-1 agonist was set to 0.60 (Armstrong, Gallagher et al. 1986).

**DR lesion/LC lesion/VTA lesion**

The Blier group did a series of experiments in male rodents where they lesioned one of the monoaminergic nuclei, then observed the change in the firing activity of the other two monoaminergic nuclei to determine how they influence one another. The results of these experiments are included in the truth table and described in detail in the supplement to the MS-model study (Camacho, Vijitbenjaronk et al. 2018).

**Stress**

The stress input sends an excitatory projection to the PVN (Sapolsky 2000, Gold and Chrousos 2002, de Kloet, Joels et al. 2005). Stress leads to maximal elevation of PVN activity as measured by induction of c-fos mRNA expression in male rats (Cullinan, Herman et al. 1995). Stress also leads to maximal increases in the activity of the adrenal gland in male rodents as measured by maximal blood flow increases after acute stress and c-fos induction (Goldman 1963, Yang, Koistinaho et al. 1989). The truth-table values for PVN and adrenal gland were set to 0.70 with stress. Stress moderately increases plasma CRF, ACTH, and cortisol in male subjects, so the truth-table values for CRF, ACTH and cortisol with stress were set to 0.60 (Kitay 1961, Viau, Bingham et al. 2005, Iwasaki-Sekino, Mano-Otagiri et al. 2009) (Goldman 1963, Zimmermann and Critchlow 1967, Harbuz and Lightman 1989, Rivier 1993). Castrated males have maximal increases in CRF, ACTH, cortisol, and AVP in response to stress, so the truth-table values for CRF, ACTH, cortisol, and AVP were all set to 0.70 for castrated males (Seale, Wood et al. 2004). When testosterone is administered to castrated males, their stress-induced increases in ACTH and cortisol return to moderate levels, so the truth-table values for ACTH and cortisol for stressed, castrated and testosterone-infused males were set to 0.60 (Handa, Nunley et al. 1994, Viau and Meaney 1996). NE levels have been found to moderately increase in response to stressful stimuli in both male and female subjects, so the truth-table value for NE with stress was set to 0.60 (Galvez, Mesches et al. 1996, Hatfield, Spanis et al. 1999). LC neuron activity in both male and female subjects has been found to moderately increase in response to stress (Abercrombie and Jacobs 1987, Buffalari and Grace 2007). Stress has also been shown to moderately increase tryptophan, 5HTP, 5HT and DA levels in many different brain regions of male subjects (Thierry, Fekete et al. 1968, Abercrombie, Keefe et al. 1989, Kawahara, Yoshida et al. 1993, Summers, Kampshoff et al. 2003). We set the truth-table values for tryptophan, 5HTP, 5HT, NE, LC and DA to 0.60. Acute stress moderately increases DR neuron firing rate in male rodents, and VTA firing rate in both male and female cats, so the truth-table values for DR and VTA were set to 0.60 (Trulson and Preussler 1984, Bambico, Nguyen et al. 2009). Acute stress also moderately increase 5HT and DA levels in castrated males, so the truth-table values for 5HT and DA with acute stress were set to 0.60 for castrated males (Doge 1993). Acute stress moderately increases glutamate levels in male rodents, so the truth-table value for glutamate was set to 0.60 (Reznikov, Grillo et al. 2007). Acute stress moderately increases AVP and oxytocin levels in male rodents, so the truth-table values for AVP and oxytocin were set to 0.60 (Hesketh, Jessop et al. 2005). The amygdala and hippocampus are moderately activated in response to stress in males while the PFC is moderately inhibited in both males and females, so the truth-table values for the amygdala, hippocampus, and PFC were set to 0.60, 0.60 and 0.40, respectively (Sakanaka, Shibasaki et al. 1986, Van de Kar, Piechowski et al. 1991, Chen, Fenoglio et al. 2006, Alexander, Hillier et al. 2007, Qin, Hermans et al. 2009). Acute stress moderately elevates melatonin levels in male rodents, so the truth-table value for melatonin was set to 0.60 (Lynch, Eng et al. 1973, Vollrath and Welker 1988). Foot-shock stress causes a moderate decrease in GABA in both male and female rats, so this truth-table value was set to 0.40 (Biggio, Corda et al. 1981). Acute immobilization-stress produces no change in galanin levels in the amygdala of male rodents, so the truth-table value for galanin with stress was set to 0.50 (Khoshbouei, Cecchi et al. 2002). Surgical stress was found to moderately decrease testosterone levels in human males, so testosterone was set to 0.50 with stress (Aono, Kurachi et al. 1976). Stress moderately increases 5HT levels in males, so the truth-table value for 5HT was set to 0.60 (Mitsushima, Yamada et al. 2006, Lanfumey, Mongeau et al. 2008, Jacobson-Pick, Audet et al. 2013). Adding progesterone to stress in males moderately increases plasma progesterone levels and prevents the rise in cortisol associated with stress (Childs, Van Dam et al. 2010). The truth-table value for progesterone and cortisol for Progesterone/Stress were set to 0.50.

**<u>Adrenalectomy, ADX/Stress, ADX/Dex and ADX/Corticosterone</u>**

With the adrenalectomy (ADX) input, the adrenal gland truth-table value was set to 0.30 (Iacobone, Albiger et al. 2008). ADX moderately increases testis activity in male rodents, so the truth-table value for testis with ADX was set to 0.60 (Desjardins and Ewing 1971). However, testosterone levels have been found to stay the same with ADX in intact male rats (Saez, Morera et al. 1977). ADX results in a maximal rise in CRF and ACTH as well as a moderate rise in AVP in both male and female subjects, so the truth table values for CRF and ACTH were set to 0.70 for AVP was set to 0.60 with adrenalectomy (Vernikos-Danellis 1965, Fink, Robinson et al. 1988,

Unno, Wu et al. 1998, Iacobone, Albiger et al. 2008). Because PVN and pituitary gland firing-rates moderately increase with ADX in male subjects, we set the truth-table values for PVN and pituitary gland to 0.60 (Kitay, Holub et al. 1959, Wynn, Harwood et al. 1985, Kasai and Yamashita 1988). ADX/Dex has been found to produce no change in testosterone levels in male rats, so the truth-table value for testosterone with ADX/Dex was set to 0.60 (Saez, Morera et al. 1977). ADX produces no change in serum LH levels in intact male rats, so the truth-table value for LH with ADX was set to 0.50 (Mann, Free et al. 1987). ADX in castrated male rats does not change LH and FSH levels, so the truth-table values for LH and FSH with ADX were set to 0.50 for castrated males (Schwartz and Justo 1977). The combination of ADX/Corticosterone was found to have no change on testis activity, but a decrease in testosterone level (Desjardins and Ewing 1971). The truth-table values for testis and testosterone were both set to 0.50 with corticosterone/ADX. The combinations of ADX/hCG, ADX/hCG/Dex, and ADX/ACTH/hCG all resulted in a moderate increase in testosterone levels in intact male rats, so testosterone was set to 0.70 for all of these combinations (Saez, Morera et al. 1977).

**Exogenous ACTH**

Exogenous ACTH projects to ACTH receptors, and raises ACTH levels, so the truth-table value for ACTH was set to 0.60 (Kitay, Holub et al. 1959). Intramuscular injections of ACTH in male horses leads to a moderate increase in cortisol levels after 2-4 hours, so the truth-table value for cortisol with exogenous ACTH was set to 0.60 (Thorn, Forsham et al. 1950, Larsson, Edqvist et al. 1979). ACTH has been shown to increase adrenal gland activity in male rats, so the truth-table value for adrenal gland with exogenous ACTH was set to 0.60 (Yang, Koistinaho et al. 1990). Exogenous ACTH for three days produces no change in LH levels but moderately decreases testosterone levels in intact male rats, so the truth-table values for LH and testosterone with exogenous ACTH were both set to 0.50 (Saez, Morera et al. 1977, Mann, Free et al. 1987). Exogenous ACTH in intact male rats has been found to decrease testis size, so the truth-table value for testis was set to 0.40 with Exogenous ACTH (Baker, Schairer et al. 1950). Exogenous ACTH has also been found to increase progesterone levels in intact male rats, so the truth-table value for progesterone with exogenous ACTH was set to 0.50 (Mann, Free et al. 1987). Exogenous ACTH moderately decreases FSH levels in intact male rats, so the truth-table value for FSH with exogenous ACTH was set to 0.40 (Mann, Free et al. 1987). Exogenous ACTH administration in castrated male rats produces no change in LH levels, so the truth-table value for LH with ACTH was set to 0.50 for castrated males (Mann, Free et al. 1987). Exogenous ACTH results in a maximal decrease in testosterone levels in castrated male rats, so the truth-table value for testosterone with ACTH was set to 0.40 for castrated males (Mann, Free et al. 1987). Exogenous ACTH results in moderately elevates cortisol and progesterone levels in castrated male rats, so the truth-table values for cortisol and progesterone were set to 0.60 and 0.50, respectively, for castrated males (Mann, Free et al. 1987). Exogenous ACTH moderately decreases FSH levels in castrated males, so the truth-table value for FSH with exogenous ACTH was set to 0.40 for castrated males (Mann, Free et al. 1987).

**Exogenous CRF**

Exogenous CRF projects to CRF1 and CRF2 receptors and elevates CRF levels in both males and females (Merchenthaler 1984). Exogenous CRF administration in both male and female humans has been found to moderately increase plasma ACTH and cortisol levels (Hermus, Pieters et al. 1984). The truth-table values for ACTH and cortisol with exogenous CRF were both set to 0.60.

**PVN, amygdala, hippocampus, and PFC lesion**

Lesions to the PVN, amygdala, hippocampus and PFC all result in maximal decreases in PVN, amygdala, hippocampus, and PFC, respectively (Chang, Tran et al. 1980). The truth-table values for PVN, amygdala, hippocampus, and PFC, with PVN lesion, amygdala lesion, hippocampus lesion, and PFC lesion, respectively, were all set to 0.30. PVN lesion maximally reduces CRF levels and moderately reduces oxytocin levels in both male and female rodents, so the truth-table values for CRF and oxytocin with PVN lesion were set to 0.30 and 0.40, respectively (Bruhn, Plotsky et al. 1984, Antoni, Fink et al. 1990). Basal ACTH levels have also been found to moderately decrease with a PVN lesion in both male and female rodents, so the truth-table value for ACTH was set to 0.40 (Makara, Stark et al. 1981). PVN lesion has been found to maximally decrease melatonin levels in male rodents, so the truth-table value for melatonin with PVN lesion was set to 0.30 (Klein, Smoot et al. 1983). Adrenal gland activity has been found to moderately decrease with PVN lesion in both male and female rodents, so the truth-table value for adrenal gland was set to 0.40 (Makara, Stark et al. 1981).

**PVN, amygdala, hippocampus, and PFC lesion and Stress**

The combination of PVN lesion and stress results in a moderate increase in CRF, ACTH, and cortisol in both males and females (Makara, Stark et al. 1981, Bruhn, Plotsky et al. 1984, Makara 1992). The truth-table values for these were set to 0.60. Amygdala lesion and stress has been shown to lead to a moderate increase in CRF, ACTH, and cortisol in male subjects, so the truth table values for these were also set to 0.60 (Sakanaka, Shibasaki et al. 1986, Van de Kar, Piechowski et al. 1991, Feldman, Conforti et al. 1994). However, lesions of the PFC and hippocampus combined with stress lead to maximal elevations of ACTH and cortisol in male subjects (Jacobs, Wise et al. 1974, Herman, Schafer et al. 1989, Jacobson and Sapolsky 1991, Diorio, Viau et al. 1993, Herman, Cullinan et al. 1995). We set the truth-table values for ACTH and cortisol with PFC lesion/Stress and Hippocampus lesion/stress to 0.70.

**PVN lesion/Gepirone**

The combination of PVN lesion and a 5HT1A receptor agonist prevented the rise in ACTH observed with the agonist by itself in male rats, so the truth-table value for ACTH with PVN lesion/Gepirone was set to 0.50 (Bluet Pajot, Mounier et al. 1995).

**Dexamethasone/CRF**

The Dexamethasone/CRF test is widely used in depression research to detect HPA axis dysfunction (Rush, Giles et al. 1996, Kunugi, Ida et al. 2006, Heim, Mletzko et al. 2008). CRF is injected after pretreatment with Dexamethasone (Kunugi, Ida et al. 2006). In normal control subjects, ACTH and cortisol secretion does not increase, but in stressed subjects, this test shows moderate increases in ACTH and cortisol secretion (Hohnloser, Von Werder et al. 1989, Kunugi, Ida et al. 2006, Heim, Mletzko et al. 2008). This was found for both males and females. We therefore set the truth-table values for ACTH and cortisol with Dexamethasone/CRF to 0.50, and the truth-table values for ACTH and cortisol with Dexamethasone/CRF/stress to 0.60.

**SSRI/Stress**

Acute restraint stress with acute SSRI has been found to moderately increase AVP levels in male rodents, so the truth-table value for AVP with SSRI/stress was set to 0.60 (Hesketh, Jessop et al. 2005). Oxytocin and CRF levels moderately increase with SSRI/stress in male rodents, so the truth-table values for oxytocin and CRF with SSRI/Stress were set to 0.60 (Hesketh, Jessop et al. 2005). The combination of SSRI/Stress maximally increases ACTH and cortisol in male rodents, so the truth-table values for ACTH and cortisol were set to 0.70 with SSRI/Stress (Hesketh, Jessop et al. 2005). SSRI/Stress has been found to maximally increase NE levels in male rodents, so the truth-table value for NE with SSRI/stress was set to 0.70 (Page and Abercrombie 1997).

**SSRI/WAY-100635**

The combination of an SSRI and a 5HT1AR antagonist (SSRI/WAY) maximally increases 5HT levels in male rodents, so the truth-table value for SSRI/WAY was set to 0.70 (Arborelius, Nomikos et al. 1996). The combination of an SSRI and WAY produces no change in DR firing rate from baseline in male and female rodents, so the truth-table value for DR with SSRI/WAY was set to 0.50 (Arborelius, Nomikos et al. 1995, Hajos, Gartside et al. 1995) (Evrard, Laporte et al. 1999)

**SSRI/Bupropion**

Co-administration of an SSRI and Bupropion (SSRI/Bupropion) has been found by the Blier group to double DR firing rate and decrease LC firing rate by 60% in male rodents (Ghanbari, El Mansari et al. 2010). The truth-table values for DR and LC with SSRI/Bupropion were set to 0.60 and 0.40, respectively. SSRI/Bupropion have been found to moderately increase 5HT, NE and DA levels in male rodents, so the truth-table values for 5HT, NE and DA with acute SSRI/Bupropion were set to 0.60 (Li, Perry et al. 2002).

**SSRI/Aripiprazole**

The Blier group found that the combination of an SSRI and Aripiprazole (SSRI/Aripiprazole) produces no significant change in the firing rates of DR or VTA neurons, but decreases LC neuron firing by 26% in male rodents (Chernoloz, El Mansari et al. 2009). The truth table values for DR, LC, and VTA with SSRI/Aripiprazole were set to 0.50, 0.40, and 0.50, respectively.

**SSRI/Quetiapine**

The Blier group found that the combination of an SSRI and Quetiapine (SSRI/Quetiapine) produces a 65% decrease in DR neuron firing and a 27% increase in LC neuron firing in male rodents (Chernoloz, El Mansari et al. 2012). The truth table values for DR and LC with SSRI/Quetiapine were therefore set to 0.40 and 0.60, respectively.

**Reboxetine/Stress**

The combination of Reboxetine and fearful stimuli (fearful faces) has been found to moderately increase amygdala activity in males and females in fMRI experiments (Onur, Walter et al. 2009). The truth-table value for amygdala with Reboxetine/Stress was set to 0.60. The combination of Reboxetine and stress in male rodents has been found to produce no change in 5HT levels, maximally increases NE levels, and moderately increase DA levels (Page and Lucki 2002). The truth-table values for 5HT, NE and DA with Reboxetine/Stress were set to 0.50, 0.70, and 0.60, respectively.

**Estrogen exogenous**

Estrogen is a hormone input that projects to alpha and beta estrogen receptors in the model (Shughrue, Lane et al. 1997). Estrogen administration in both male and female intact rats has been shown to moderately increase DR neuron activity, so the truth-table value for DR was set to 0.60 for exogenous estrogen (Robichaud and Debonnel 2005). Estrogen produces no change CRF or AVP levels in castrated male rats, so the truth-table values for CRF and AVP with estrogen were set to 0.50 for castrated males (Patchev, Hayashi et al. 1995).

**Testosterone exogenous**

Testosterone projects to the testosterone receptor in the model. Acute (3-day) administration of testosterone moderately increases DR neuron firing in both male and freely cycling female rats, so the truth-table value for DR with testosterone was set to 0.60 (Robichaud and Debonnel 2005). Testosterone administration to male rats was found to moderately increase 5HT and DA levels, so the truth-table values for 5HT and DA were set to 0.60 (de Souza Silva, Mattern et al. 2009).

**hCG**

Administration of hCG to intact male rats has been shown to moderately increase testosterone levels, so the truth-table value for testosterone was set to 0.70 (Saez, Morera et al. 1977). The combination of hCG and Dexamethasone as well as the combination of hCG and Corticosteroid to male testicular cell culture were both found to moderately decrease testosterone levels, so the truth-table values for testosterone with hCG/Dex and hCG/Corticosteroid were set to 0.50 (Bambino and Hsueh 1981).

**ER-beta agonist**

ER-beta agonist projects to the Beta estrogen receptor. Administration of the ER-beta agonist to intact male rodents produces no change in testosterone levels, so the truth-table value for testosterone with ER-beta agonist was set to 0.60 (Patisaul, Burke et al. 2009). ER-beta agonist moderately increases 5HT and DA levels after 3-hours of administration in male rodents, so the truth-table values for 5HT and DA were set to 0.60 with acute ER-beta agonist (Hughes, Liu et al. 2008).

**ER-alpha agonist**

ER-alpha agonist projects to the Alpha estrogen receptor. Administration of the ER-alpha agonist to intact male rodents produces no change in testosterone levels, so the truth-table value for testosterone with ER-alpha agonist was set to 0.60 (Patisaul, Burke et al. 2009).

**AVP Exogenous**

Exogenous AVP binds to V1A receptors in the model. Exogenous AVP has been found to moderately increase ACTH and cortisol levels in male humans, so the truth-table values for ACTH and cortisol with Exogenous AVP were set to 0.60 (Hensen, Hader et al. 1988).

**GnRH Exogenous**

Exogenous GnRH is a hormone input that binds to the GnRH receptor. Exogenous GnRN administration for three days to intact male rats moderately increases LH, FSH, and testosterone levels, so the truth-table values for LH, FSH and testosterone with exogenous GnRH were set to 0.60, 0.60, and 0.70 (Bruni, Huang et al. 1977, Mann, Free et al. 1987).

**ACTH Exogenous/GnRH Exogenous**

The combination of Exogenous ACTH and GnRH resulted in no change in serum testosterone levels in intact male rats, so the truth-table value for testosterone with Exogenous ACTH/GnRH were set to 0.60 (Mann, Free et al. 1987). The combination of exogenous ACTH and GnRH resulted in a moderate increase in LH levels in intact male rats, so the truth-table value for LH with exogenous ACTH/GnRH was set to 0.60 (Mann, Free et al. 1987).

**ADX/ACTH Exogenous/GnRH Exogenous**

The combination of ADX, exogenous ACTH and exogenous GnRH produces a moderate increase in LH levels in intact male rats, so the truth table value for LH with ADX/ACTH/GnRH was set to 0.60 (Mann, Free et al. 1987).

**ADX/ACTH Exogenous**

The combination of ADX and exogenous ACTH produces no change in testosterone levels of intact male rats, so the truth-table value of testosterone with ADX/ACTH was set to 0.60 (Saez, Morera et al. 1977, Mann, Free et al. 1987).

**ADX/GnRH**

The combination of ADX/GnRH produces no change in testosterone levels in intact male rats, so the truth-table value for testosterone with ADX/GnRH was set to 0.60 (Mann, Free et al. 1987).

**Corticosterone**

Corticosterone is a hormone input that projects to MC and GC receptors in the model. Corticosterone treatment moderately decreases adrenal gland weight in male rats, so the truth-table value for adrenal gland was set to 0.40 with corticosterone (Mann, Free et al. 1987). Serum LH levels were found to be unaffected by corticosterone in male rats, so the truth-table value for LH was set to 0.50 with corticosterone (Mann, Free et al. 1987). Testosterone levels were found to moderately decrease with acute corticosterone in male rats, so the truth-table value for testosterone with corticosterone was set to 0.50 (Mann, Free et al. 1987).

**Cortisol Infusion**

Cortisol infusion in castrated male Rhesus monkeys resulted in a maximal rise in cortisol levels, so the truth-table value for cortisol was set to 0.70 for cortisol infusion in castrated males. Cortisol infusion in castrated male Rhesus monkeys moderately decreases FSH and LH levels, but the addition of GnRH to the Cortisol infusion in castrated males results in in normal FSH and LH levels. The truth-table values for FSH and LH for cortisol infusion in castrated males were set to 0.40 and the truth-table values for FSH and LH for cortisol infusion in castrated males and GnRH were set to 0.50 (Dubey and Plant 1985).

**Corticosterone/GnRH**

The combination of corticosterone/GnRH moderately increases LH levels, does not change testosterone levels or testes size, and decreases adrenal gland size in male rats (Mann, Free et al. 1987). The truth-table values for LH, testosterone, testes, and adrenal gland were set to 0.60, 0.50, 0.50, and 0.40, respectively, with corticosterone/GnRH.

**Castration**

Castration has an inhibitory projection to the testes in the model. Castration has been found to maximally decrease testosterone levels and produce no change in DR neuron firing rate, so the truth-table value for testosterone was set to 0.40 and for DR was set to 0.50 (Robichaud and Debonnel 2005). Castration in intact male rats has also been found to moderately increase LH, FSH and progesterone levels, so the truth-table values for LH, FSH and progesterone with castration were set to 0.60, 0.60, and 0.50, respectively (Schwartz and Justo 1977, Mann, Free et al. 1987).

**5HT exogenous**

Exogenous 5HT injections in intact male rats increases 5HT levels and decreases testis weight, Gepso the truth-table value for 5HT was set to 0.60 and for testis was set to 0.40 with acute 5HT exogenous. Exogenous 5HT also increases LH and FSH levels, but decreases testosterone levels. The truth-table values for LH and FSH were set to 0.60, and for testosterone was set to 0.50 for exogenous 5HT (Hedger, Khatab et al. 1995).

**S3: Hardware Considerations**

All LTL analyses were conducted in Python™ using an Intel Core i5 processor with 4, 2.80 GHz cores with 8.00 GB of RAM. All other analyses were carried out in MATLAB® using an Intel Core 2 Duo CPU processors with 2, 2.33 GHz cores and 4.00 GB of RAM under the Windows 7 operating system, an Intel-inside CORE i7 processor with 2, 2.69 GHz cores and 8.00 GB of RAM under the Windows 8 operating system, and an Intel Core i7 processor with 4, 4.00 GHz cores and 32.00 GB of RAM under the Windows 10 operating system. Limitations in computer memory prevented computing the full set of adjustable-TSC strength configurations out to 7 adjustment steps.

**S4: Setting Therapeutic Criteria**

Therapeutic criteria were set using the same criteria as described in the MS-model (Camacho, Vijitbenjaronk et al. 2018). Briefly, chronic SSRI administration has been shown to double 5HT levels associated with acute SSRI administration (Ceglia, Acconcia et al. 2004). The desired baseline 5HT level was set to 0.50, and the desired 5HT output for acute SSRI was set to 0.60. The therapeutic 5HT floor was therefore set to 0.70 in order to double the increase of 0.10 observed with acute SSRI. For consistency, 0.70 was also used as the therapeutic floor for NE and DA although therapeutic NE and DA levels resulting from chronic antidepressant administration have not been determined. Because the high CORT levels associated with acute SSRI have been found to decrease with chronic SSRI (Ruhe, Khoenkhoen et al. 2015), the therapeutic ceiling for CORT was set to 0.70. This is the same level assigned to CORT in the truth table due to acute SSRI, reflecting experimental observations (see S2 for SSRI above).

**S5: Details on LTL analysis**

The following predicates were used in the LTL analysis: `fht_high`, 5HT is above the 5HT therapeutic floor (>= 0.70); `cort_low`, CORT is below the therapeutic CORT ceiling (<=0.70); `TSC_sens_gt_3`, the adjustable TSC has sensitized by at least 3 steps; `TSC_desens_gt_3`, the adjustable TSC has desensitized by at least 3 steps, where TSC is a placeholder for any 1 of the 13 adjustable TSCs; and `Adapted`, the state has adaptation error at least 25% less than initial error. Then we evaluated the following propositions for each of the 13 adjustable TSCs, where `|->`, `/\`, `\/`, and ~ are the LTL operators for "LEADS TO", "AND", "OR", and "NOT", respectively:

```
TSC_sens_gt_3 |-> (Adapted /\ fht_high /\ cort_low) \/
~TSC_sens_gt_3

TSC_desens_gt_3 |-> (Adapted /\ fht_high /\ cort_low) \/
~TSC_desens_gt_3
```

If there are single or few TSCs determinative of the therapeutic state, then large subtrees of the state transition tree should be either uniformly adapted and therapeutic, or uniformly not adapted and/or non-therapeutic, depending on the polarity of those few TSCs. We attempt to determine the extent to which transitions from adapted and therapeutic TSC-strength configurations (i.e. reference states) lead to configurations which are also adapted and therapeutic. Since this question deals with transitions between pairs of states reachable from one another using incremental adjustments, it can be framed in the context of LTL using the NEXT operator O and the generic predicate P standing here for states that are adapted and therapeutic:

```
P = Adapted /\ fht_high /\ cort_low
```

We then evaluated `O(P)`, which takes a predicate P and returns True if and only if that predicate holds at the state reached after one transition from the reference state. This operator can be nested; the predicate `O(O(P))` returns True if and only if P holds at the state reached after two transitions, and so on for greater numbers of transitions from the reference state. Then the proposition:

```
P /\ O(P)
```

can be used to find all degree-1 (i.e. 1 adjustment) transitions from an adapted/therapeutic reference state to a subsequent adapted/therapeutic state. This may be extended to higher degrees in a straightforward manner. For example, the proposition:

```
P /\ O(P) /\ O(O(P))
```

can be used to find all degree-2 (i.e. 2 adjustment) sequences from an adapted/therapeutic reference state to a subsequent adapted/therapeutic state and on to another subsequent adapted/therapeutic state. This proposition corresponds with the intuitive notion that the reference state itself is adapted and therapeutic, the state reachable in one transition is adapted and therapeutic, and the state reachable in two transitions is also adapted and therapeutic. The extension to higher-degree sequences is straightforward. We searched for sequences of all degrees starting from reference, adapted/therapeutic TSC-strength configurations and proceeding up to degree-6. Since the state transition tree included many duplicate states and sequences all duplicates were removed before statistics were computed. Proportions of adapted and therapeutic sequences of various degrees reported in Results as percentages of the total number of unique adapted and therapeutic states.

**Supplemental Figure 1: Complete model structure schematic.** This diagram shows all model structure connections. Green, red and blue lines represent excitatory, inhibitory, and trained (determined by the training algorithm) valences, respectively, between model elements (units). Unit type (input, receptor, transmitter/hormone, or precursor/metabolite/enzyme/transporter) is represented by a different shape. All units of the same type are in the same row. The file CompleteMSSModelStructure.jpeg, which is included in the Supplementary Material for this paper, contains this diagram and can be viewed interactively.

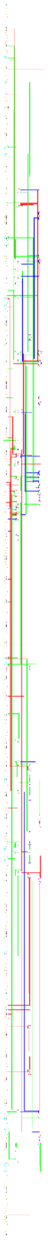

**Supplemental Figure 2: Close agreement between desired (i.e. target) and actual output responses after training but before pruning (A−C) and after pruning and re-training (D−F).** The colormap scales for the desired and actual output plots (A,B,C,D, and E) range from 0.00 to 0.70. Each row represents a single input pattern and each column represents an output value. The first row represents baseline output values when no inputs are present. The colormap for the absolute differences between the desired and actual output values (C and F) are between 0.00 and 4.50x10$^{-3}$ for (C) and 0.00 and 3.50x10$^{-3}$ for (F). The networks were pruned to minimize non-structure connections. The RMS error over all training patterns was 2.02x10$^{-4}$ for the unpruned network in (A-C) and 1.84x10$^{-4}$ for the pruned network in D-F.

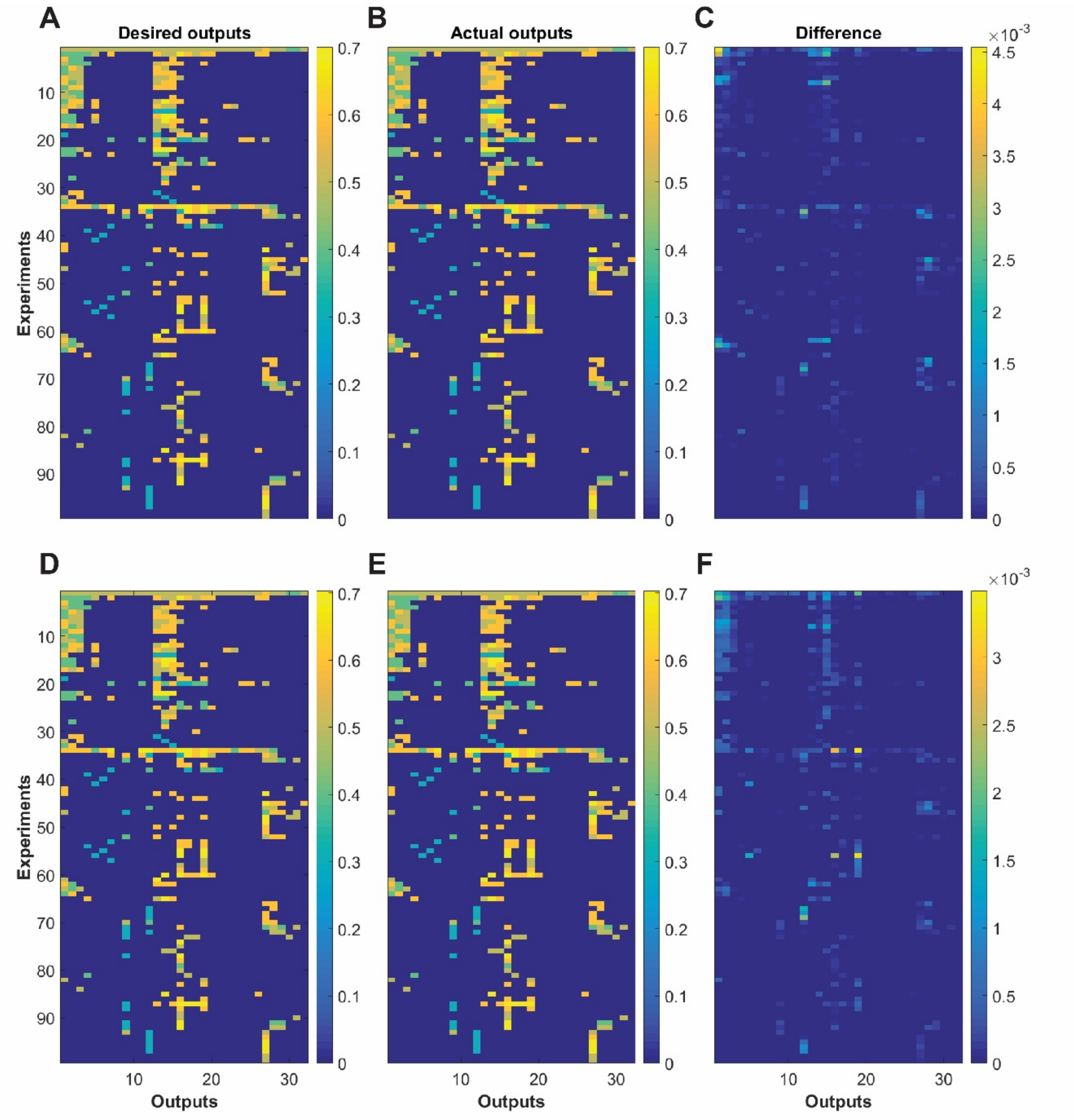

**Supplemental Table 1: Complete MSS-model truth-table.** The complete model truth-table is available as a spreadsheet in file CompleteMSSModelTruthTable.xls, which is included with the Supplemental Material for this paper. Each row of the truth table in represents observed input-output relationships derived from human or animal experiments on the monoaminergic, stress hormone, and sex hormone systems. The inputs include drugs, hormones, lesions, or combinations thereof. Inputs are either present (1) or absent (0). Most rows incorporate findings compiled from multiple experiments, sometimes from multiple labs. These findings are presented to the model as input/desired-output patterns during the training procedure. Abbreviations: amygdala, AMY; prefrontal cortex, PFC; hippocampus, HIPP; paraventricular nucleus of the hypothalamus, PVN; preoptic area, POA; cortisol, CORT; cortocotropin-releasing factor, CRF; arginine vasopressin, AVP; estrogen receptor, ER; alpha-1R, α-1 adrenergic receptor; CRF1 receptor, CRF1R; human chorionic gonadotropin, hCG; gonadotropin releasing hormone, GnRH; adrenocorticotropin hormone, ACTH; gamma-Aminobutyric acid, GABA; tryptophan, Trp; 5-hydroxytryptophan, 5HTP; selective serotonin reuptake inhibitor, SSRI.

**Supplemental Table 2: Canonical MSS-model weights.** The MSS-model has 36 weights classified as "canonical weights," because these weights mediate the known interactions of the monoaminergic nervous system, the stress-steroid system, and the sex-steroid system. The lower bound of the canonical weights was set to |1| during training. Upon further analysis it was determined that the canonical weights developed the strongest weights in the networks, and each network was most sensitive to changes in the canonical weights. The pruning procedure further increased network sensitivity to the canonical weights.

| Number | Weight | Polarity | References |
|---|---|---|---|
| 1 | DR to 5HT | + | (Hensler, Ferry et al. 1994, Monti 2010) |
| 2 | 5HT to 5HT1AR | + | (Davidson and Stamford 1995, Blier, Pineyro et al. 1998) |
| 3 | 5HT1AR to DR | − | (Davidson and Stamford 1995, Azmitia, Gannon et al. 1996) |
| 4 | 5HTT to 5HT | − | (Lesch, Aulakh et al. 1993, Hensler, Ferry et al. 1994) |
| 5 | LC to NE | + | (Grenhoff, Nisell et al. 1993, Samuels and Szabadi 2008) |
| 6 | NE to AR2 | + | (Cedarbaum and Aghajanian 1977, Washburn and Moises 1989) |
| 7 | AR2 to LC | − | (Cedarbaum and Aghajanian 1977, Washburn and Moises 1989) |
| 8 | NET to NE | − | (Iversen 2000, Bonisch and Bruss 2006) |
| 9 | VTA to DA | + | (Ornstein, Milon et al. 1987, Guiard, El Mansari et al. 2008) |
| 10 | DA to D2R | + | (Benoit-Marand, Borrelli et al. 2001, Perra, Clements et al. 2011) |
| 11 | D2R to VTA | − | (Hall, Sedvall et al. 1994, Perra, Clements et al. 2011, Koyama, Mori et al. 2014) |
| 12 | DAT to DA | − | (Ciliax, Heilman et al. 1995, Iversen 2000) |
| 13 | PVN to CRF | + | (Makara, Stark et al. 1981, Bruhn, Plotsky et al. 1984) |
| 14 | CRF to CRF1R | + | (Van Pett, Viau et al. 2000, Hauger, Risbrough et al. 2006, Holsboer and Ising 2008) |
| 15 | CRF1R to Corticotroph | + | (Van Pett, Viau et al. 2000, Nikodemova, Diehl et al. 2002) |
| 16 | Corticotroph to ACTH | + | (Makara, Stark et al. 1981, Bruhn, Plotsky et al. 1984) |
| 17 | ACTH to ACTHR | + | (Xia and Wikberg 1996, Papadimitriou and Priftis 2009) |
| 18 | ACTHR to Adrenal Gland | + | (Yang, Koistinaho et al. 1990, Xia and Wikberg 1996, Papadimitriou and Priftis 2009) |

| | | | |
|---|---|---|---|
| 19 | Adrenal Gland to CORT | + | (Grant, Forrest et al. 1957, Papadimitriou and Priftis 2009) |
| 20 | CORT to GCR | + | (Pariante and Miller 2001, Papadimitriou and Priftis 2009) |
| 21 | GCR to Adrenal Gland | − | (Loose, Do et al. 1980, Kalinyak, Dorin et al. 1987) |
| 22 | GCR to Corticotroph | − | (Morimoto, Morita et al. 1996, Ozawa, Ito et al. 1999) |
| 23 | GCR to PVN | − | (Morimoto, Morita et al. 1996, Ozawa, Ito et al. 1999) |
| 24 | POA to GnRH | + | (Pfaus, Jakob et al. 1994) |
| 25 | GnRH to GnRHR | + | (Kakar, Musgrove et al. 1992) |
| 26 | GnRHR to Gonadotroph | + | (Shacham, Harris et al. 2001) |
| 27 | Gonadotroph to FSH | + | (Shacham, Harris et al. 2001) |
| 28 | Gonadotroph to LH | + | (Shacham, Harris et al. 2001) |
| 29 | FSH to FSHR | + | (Dierich, Sairam et al. 1998) |
| 30 | LH to LHR | + | (Jia, Oikawa et al. 1991) |
| 31 | FSHR to Testes | + | (Catt, Baukal et al. 1979, Heckert and Griswold 1991) |
| 32 | LHR to Testes | + | (Belanger, Auclair et al. 1979, Catt, Baukal et al. 1979) |
| 33 | Testes to Testosterone | + | (Royland, Weber et al. 1994) |
| 34 | Testosterone to AR | + | (Grino, Griffin et al. 1990) |
| 35 | AR to POA | − | (Handa, Kerr et al. 1996) |
| 36 | AR to Gonadotroph | − | (Herbison, Skinner et al. 1996) |

**Supplemental Table 3: Adjustable MSS-model TSCs.** A subset of the model transmitter-system components (TSCs) are known to be adaptable under the conditions of the experiments from which the truth table was derived. These 13 TSCs are referred to as “adjustable TSCs” and are comprised of canonical weights from each of the 3 systems (monoaminergic, stress-steroid, sex-steroid) in the model. Adjusted TSC configurations that reduced adaptation error by bringing responses of the monoaminergic brain regions (DR, LC, and VTA) back toward their baseline values were considered adapted.

| Number | Weight | Polarity | References |
|---|---|---|---|
| 1 | 5HT1AR to DR | − | (Blier and de Montigny 1987, Szabo and Blier 2001, El Mansari, Ghanbari et al. 2008, Ghanbari, El Mansari et al. 2010, Rozeske, Evans et al. 2011) |
| 2 | AR2 to LC | − | (Szabo and Blier 2002, El Mansari, Ghanbari et al. 2008) |
| 3 | D2R to VTA | − | (Chernoloz, El Mansari et al. 2009, Katz, Guiard et al. 2010, Madhavan, Argilli et al. 2013) |
| 4 | 5HTT to 5HT | − | (Lesch, Aulakh et al. 1993, Benmansour, Cecchi et al. 1999, Lau, Horschitz et al. 2008) |
| 5 | NET to NE | − | (Hebert, Habimana et al. 2001, Miner, Jedema et al. 2006, Pietrzak, Gallezot et al. 2013) |

| | | | |
|---|---|---|---|
| 6 | DAT to DA | − | (Neumeister, Willeit et al. 2001, Brunswick, Amsterdam et al. 2003, Kugaya, Seneca et al. 2003, Yang, Yeh et al. 2008) |
| 7 | GCR to PVN | − | (Pariante and Miller 2001, Barden 2004, Ladd, Huot et al. 2004) |
| 8 | GCR to Corticotroph | − | (Barden 2004, Ladd, Huot et al. 2004) |
| 9 | GCR to Adrenal Gland | − | (Barden 2004, Ladd, Huot et al. 2004) |
| 10 | AR to POA | − | (Handa, Kerr et al. 1996) |
| 11 | AR to Gonadotroph | − | (Bremner, Millar et al. 1994) |
| 12 | FSHR to Testes | + | (Catt, Baukal et al. 1979, Heckert and Griswold 1991) |
| 13 | LHR to Testes | + | (Belanger, Auclair et al. 1979, Catt, Baukal et al. 1979) |
